# Inhibition mechanism and antiviral activity of an α-ketoamide based SARS-CoV-2 main protease inhibitor

**DOI:** 10.1101/2023.03.09.531862

**Authors:** Xiaoxin Chen, Xiaodong Huang, Qinhai Ma, Petr Kuzmič, Biao Zhou, Jinxin Xu, Bin Liu, Haiming Jiang, Wenjie Zhang, Chunguang Yang, Shiguan Wu, Jianzhou Huang, Haijun Li, Chaofeng Long, Xin Zhao, Hongrui Xu, Yanan Sheng, Yaoting Guo, Chuanying Niu, Lu Xue, Yong Xu, Jinsong Liu, Tianyu Zhang, James Spencer, Wenbin Deng, Shu-Hui Chen, Xiaoli Xiong, Zifeng Yang, Nanshan Zhong

**Author notes:** These authors contributed equally: Xiaoxin Chen, Xiaodong Huang, Qinhai Ma, Petr Kuzmič.

## Abstract

SARS-CoV-2 has demonstrated extraordinary ability to evade antibody immunity by antigenic drift. Small molecule drugs may provide effective therapy while being part of a solution to circumvent SARS-CoV-2 immune escape. In this study we report an α-ketoamide based peptidomimetic inhibitor of SARS-CoV-2 main protease (M^pro^), RAY1216. Enzyme inhibition kinetic analysis established that RAY1216 is a slow-tight inhibitor with a *K*i of 8.6 nM; RAY1216 has a drug-target residence time of 104 min compared to 9 min of PF-07321332 (nirmatrelvir), the antiviral component in Paxlovid, suggesting that RAY1216 is approximately 12 times slower to dissociate from the protease-inhibitor complex compared to PF-07321332. Crystal structure of SARS-CoV-2 M^pro^:RAY1216 complex demonstrates that RAY1216 is covalently attached to the catalytic Cys145 through the α-ketoamide warhead; more extensive interactions are identified between bound RAY1216 and M^pro^ active site compared to PF-07321332, consistent with a more stable acyl-enzyme inhibition complex for RAY1216. In cell culture and human ACE2 transgenic mouse models, RAY1216 demonstrates comparable antiviral activities towards different SARS-CoV-2 virus variants compared to PF-07321332. Improvement in pharmacokinetics has been observed for RAY1216 over PF-07321332 in various animal models, which may allow RAY1216 to be used without ritonavir. RAY1216 is currently undergoing phase III clinical trials (https://clinicaltrials.gov/ct2/show/NCT05620160) to test real-world therapeutic efficacy against COVID-19.

## Introduction

SARS-CoV-2 has become established in the human population through the coronavirus disease 2019 (COVID-19) pandemic and is likely to remain in circulation. Owing to multinational efforts, vaccines were rapidly rolled out in the early stage of the pandemic and proved successful in saving lives. However, likely due to population immune pressures established by infections and vaccinations, SARS-CoV-2 Omicron variants with highly mutated spike proteins quickly emerged (Tian et al., 2022). Rapid emergence of highly mutated variants has demonstrated the virus’s extraordinary capacity to escape humoral immunity, representing a great challenge to vaccines and therapeutic antibodies (Cox et al., 2022; Harvey et al., 2021).

A number of small molecule SARS-CoV-2 therapeutics have been developed (Fenton and Keam, 2022). This therapeutic strategy may be part of a solution to combat SARS-CoV-2 immune escape. Of note, the orally available drugs molnupiravir and Paxlovid have been approved for COVID-19 treatment after being validated through clinical trials. Molnupiravir (LAGEVRIO, also known as EIDD-2801) is a prodrug of N-hydroxycytidine; this mutagenic ribonucleoside is a broad-spectrum antiviral agent targeting the viral RNA polymerase by lethal mutagenesis. However, this molecule has also been shown to be mutagenic to the host (Zhou et al., 2021). Paxlovid is a combination of PF-07321332 (nirmatrelvir) and ritonavir. PF-07321332 is a peptidomimetic that selectively inhibits the SARS-CoV-2 main protease (M^pro^, also known as 3C-like (3CL) protease) (Owen et al., 2021; Zhao et al., 2022), while ritonavir is a cytochrome P450 inhibitor that functions to slow down cytochrome P450-mediated metabolism of PF-07321332 to improve bioavailability. However, the usage of ritonavir limits the clinical application range of Paxlovid due to the drug-drug interaction, which may cause potential safety issues. Therefore, our original goal is to aim for a drug candidate endowed with a longer half-life while maintaining good enzyme inhibitory potency as demonstrated by PF-07321332. We expect that such a newly designed M^pro^ inhibitor may possess prolonged pharmacokinetic stability in human, which can hopefully avoid the usage of ritonavir. The drug target of PF-07321332, M^pro^, plays a role in the viral polyprotein pp1a and pp1ab processing that is essential in the SARS-CoV-2 life cycle (Ziebuhr et al., 2000). The M^pro^ gene has been observed to be relatively conserved among various SARS-CoV-2 variants, therefore M^pro^ represents a promising target for drug development for SARS-CoV-2.

Other than PF-07321332, multiple series of SARS-CoV-2 M^pro^ inhibitors have been developed or discovered (Boras et al., 2021; Breidenbach et al., 2021; Dai et al., 2020; Drayman et al., 2021; Gao et al., 2022; Jin et al., 2020; Kitamura et al., 2022; Ma et al., 2020; Ma et al., 2021; Owen *et al*., 2021; Qiao et al., 2021; Quan et al., 2022; Unoh et al., 2022; Zaidman et al., 2021; Zhang et al., 2020; Zhu et al., 2020b). With a few exceptions (Breidenbach *et al*., 2021; Gao *et al*., 2022; Jin *et al*., 2020; Unoh *et al*., 2022; Zaidman *et al*., 2021), the majority of these molecules are peptidomimetics which often exhibit poor pharmacokinetic (PK) properties. In this study, we report a further peptidomimetic M^pro^ inhibitor - RAY1216 currently in phase III clinical trial. Inspired by the successful HCV protease inhibitor discovery program reported for telaprevir (Chen and Tan, 2005; Kwong et al., 2011; Yip et al., 2004a; Yip et al., 2004b), RAY1216 was developed to feature an α-ketoamide warhead and incorporates chemical moieties known to confer selectivity towards coronavirus M^pro^. Here we characterize in detail the kinetics of SARS-CoV-2 M^pro^ inhibition by RAY1216 and determine the crystal structure of the covalent adduct with SARS-CoV-2 M^pro^. Further, the antiviral activity, protection against SARS-CoV-2 variants in animal models, and PK properties are reported, and compared to those of PF-07321332.

## Structure of RAY1216

RAY1216 (**Fig. 1**) was developed via multiple rounds of optimization conducted at P1, P2, P3, and P4 moieties and finally the covalent warhead was changed from nitrile to α-ketoamide moiety. The details of the structure–activity relationship (SAR) optimizations will be further disclosed in a separate report. RAY1216 was chemically synthesized (**Fig. S1**) and the identity of the product is confirmed by NMR (**Fig. S2-S4**). The inhibitor features a cyclopentyl substituted *α*-ketoamide warhead, a pyroglutamine with a pyrrolidinone sidechain at P1 (this moiety is known to mimic glutamine, which dominates in the P1 position of coronavirus M^pro^ recognition sequences (Xiong et al., 2021)), a P2 cyclopentylproline, a P3 cyclohexylglycine and a P4 tri-fluoroacetamide (**Fig. 1**). The absolute configuration of synthesized RAY1216 was confirmed by X-ray crystallography (**Fig. S5**).

**Fig 1.**
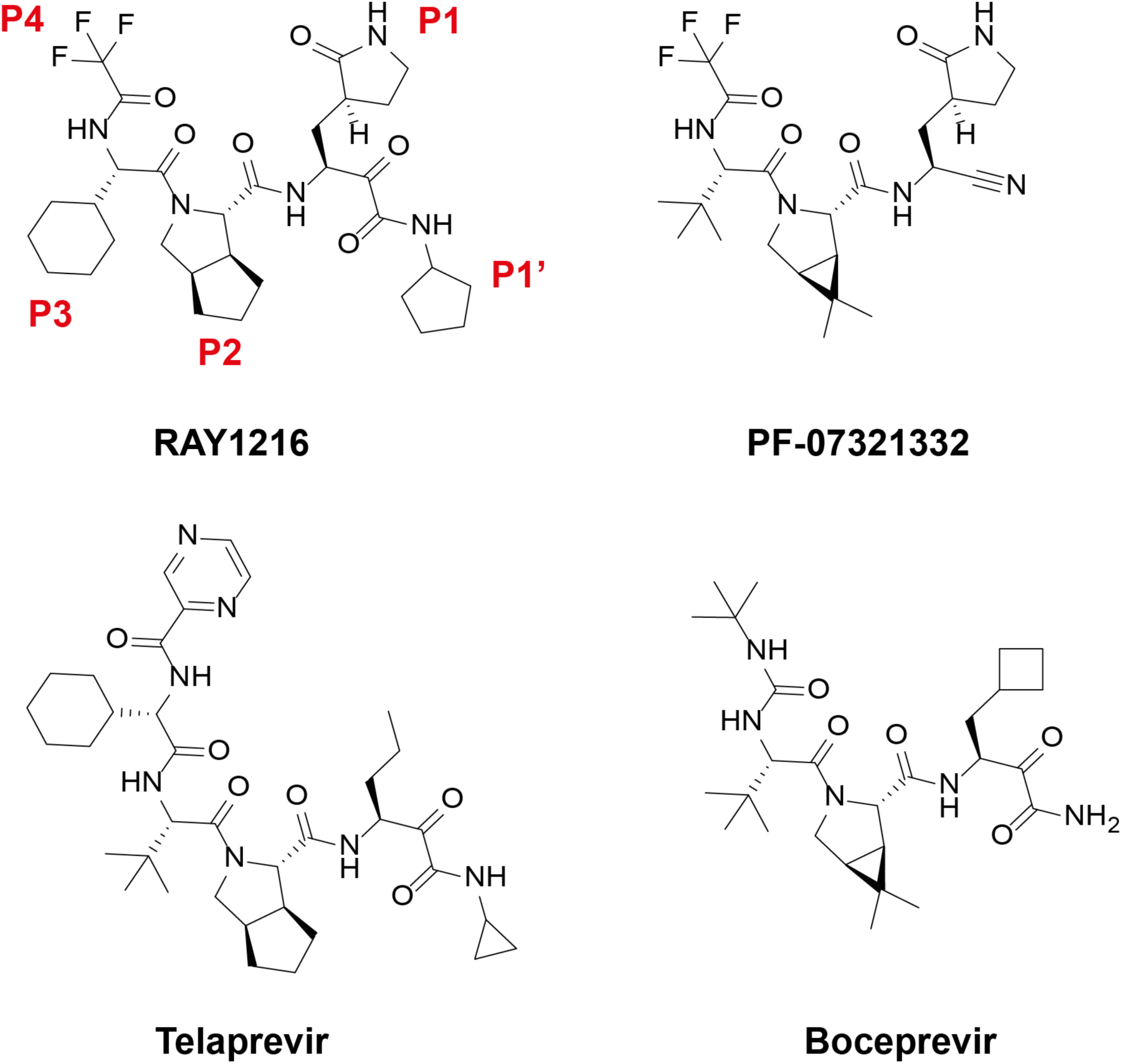
Chemical structures of RAY1216 and related anti-viral protease inhibitors.

## *In vitro* inhibition of M^pro^ by RAY1216 compared to PF-07321332

We used a fluorescence resonance energy transfer (FRET)-based peptide cleavage assay (Grum-Tokars et al., 2008) to monitor SARS-CoV-2 M^pro^ activity (**Fig. S6-7**) and we estimated a *K*_M_ of 31 μM and a *k*_cat_ of 0.12 s^−1^ for M^pro^ (**Fig. S6-8** and **Table S2-S3**). To compare inhibition by RAY1216 and PF-07321332, M^pro^ (final concentration 80 nM as determined by the Bradford assay) was added to a solution of substrate (20 μM) and inhibitor (maximum concentration 444 nM, 2:3 dilution series down to 17 nM) in the assay buffer. The increase in fluorescence intensity was monitored in real time over a period of one hour. Representative replicates for RAY1216 or PF-07321332 are shown in **Fig. 2** (also see **Fig. S8 and S9**). Both compounds displayed a gradual onset of inhibitory activity; an initial relatively uninhibited phase in product formation is followed by a gradual approach to pseudo-equilibrium (“slow binding” inhibition (Morrison, 1982; Morrison and Walsh, 1988)). Compound concentrations significantly lower than the nominal enzyme concentration caused a prominent inhibitory effect (“tight binding” inhibition (Cha, 1975; 1976; Cha et al., 1975)). The time course of the assay in the absence of inhibitors ([I] = 0) was markedly nonlinear due to substrate depletion. Under these particular experimental conditions, the classic algebraic “*k*_obs_” methods of enzyme kinetic analysis (Copeland, 2013) cannot be utilized. Instead, combined progress curves obtained at various inhibitor concentrations were fit globally to a system of first-order ordinary differential equations (ODE) solved by the software package DynaFit (Kuzmic, 1996; 2009).

**Fig. 2.**
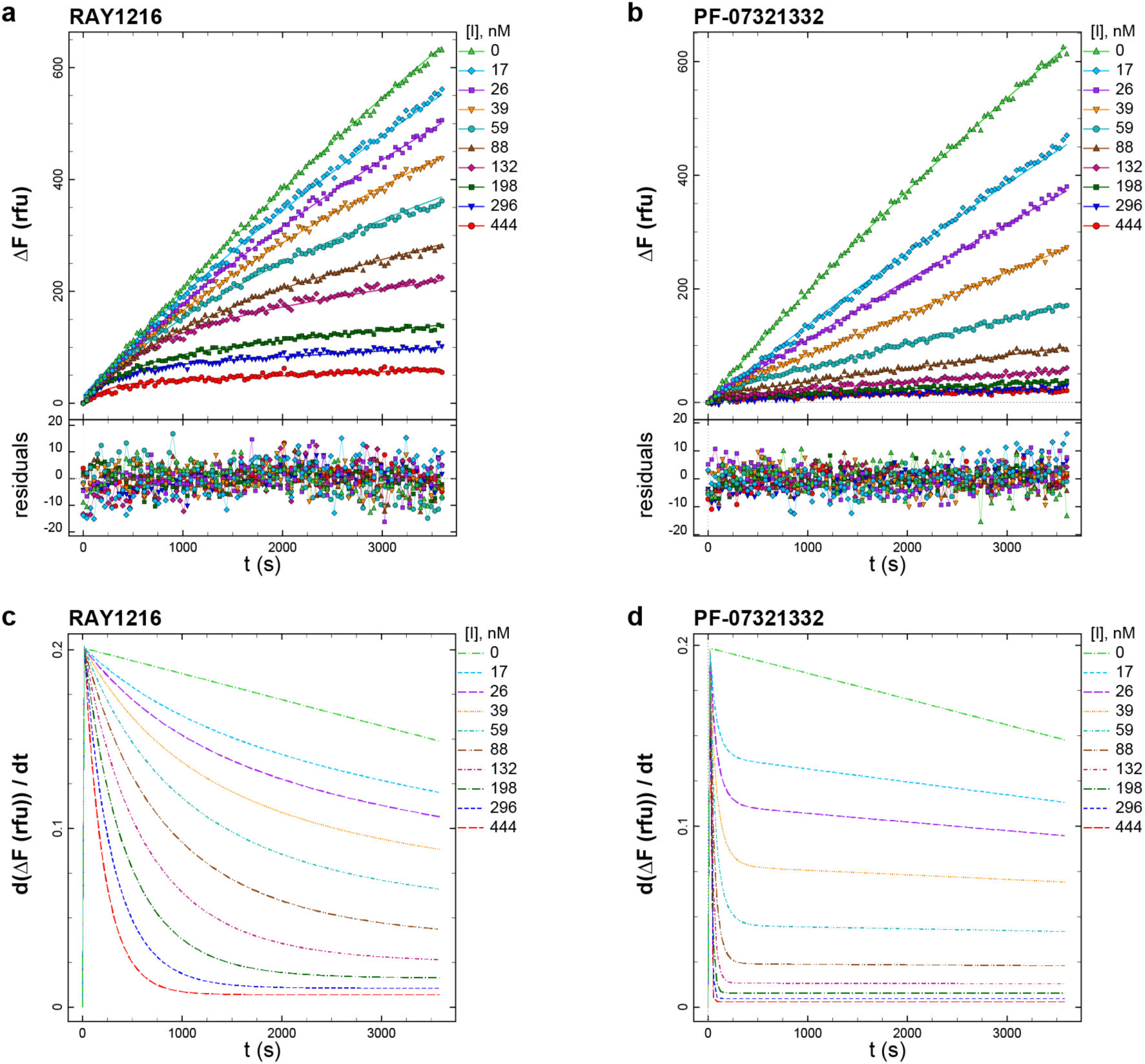
SARS-CoV-2 M^pro^ inhibition by RAY1216 and PF-07321332. **a**, Progress curves of M^pro^ inhibition (80 nM M^pro^, 20 μM substrate) at different RAY1216 concentrations (data points), reactions were started without preincubation. Progress curves are fit in DynaFit (Kuzmic, 1996; 2009) using ODE method (lines) and residuals of the fits are shown. **b**, Progress curves of M^pro^ inhibition by PF-07321332 under the same experimental conditions and they are fit in DynaFit using the same analysis procedure. **c** and **d**, Instantaneous reaction rates derived from the fits to the progress curves. See *Methods* for mathematical and statistical details of data-analytic procedures.

The data vs. model overlay plots in **Fig. 2a** and **Fig. 2b** illustrate that the overall inhibitory potencies of RAY1216 and PF-07321332 are very similar. Note that at the three highest inhibitor concentrations ([I] = 444, 296, and 198 nM) the reaction progress curves become nearly horizontal at the end of the assay both in **Fig. 2a** and **Fig. 2b**. However, also note that the approach to the quasi steady-state is markedly slower for RAY1216 when compared with PF-07321332. This fundamental difference between the two compounds is made most clearly visible in the instantaneous rate plots shown in **Fig. 2c** and **Fig. 2d**, respectively. For example, at the highest inhibitor concentration ([I] = 444 nM, bottom curve shown in red in **Fig. 2c**) it takes approximately 20 minutes for the enzyme to become fully inhibited by RAY1216. In contrast, it takes less than one minute for the enzyme to become fully inhibited by PF-07321332 under identical conditions. Note in **Fig. 2c** and **Fig. 2d** that the reaction rate does not decrease to zero even at inhibitor concentrations significantly higher than the enzyme concentration. This demonstrates the effective kinetic reversibility of the observed enzyme–inhibitor interactions despite the fact that the crystal structure shows a covalent binding mode (see below). Thus, RAY1216 appears to be an example of a “reversible covalent” inhibitor (Bradshaw et al., 2015). Since the equilibrium dissociation constants *K*_i_ = *k*_d_ / *k*_a_ for the two compounds are similar (**Table 1**), while it takes very much longer for RAY1216 to fully associate with the enzyme, it necessarily means that not only the association rate constant but also the dissociation rate constant is very much lower for RAY1216, in comparison with PF-07321332. In that sense, RAY1216 could be described as a “slow-on, slow-off” inhibitor, whereas PF-07321332 inhibition of M^pro^ is “fast-on, fast-off”.

**Table 1.** Kinetic parameters of M^Pro^ inhibition by RAY1216 and PF-07321332 as determined by ODE method in DynaFit. Mean and standard deviation from replicates (n = 3) are reported. See Methods for mathematical and statistical details of data-analytic procedures.

| parameter | unit | RAY1216 | PF-07321332 | ratio RAY/PF |
| --- | --- | --- | --- | --- |
| $k_a$ | $\mu\text{M}^{-1} \text{s}^{-1}$ | $0.019 \pm 0.001$ | $0.49 \pm 0.04$ | 0.04 |
| $k_d$ | $\text{s}^{-1}$ | $0.000161 \pm 0.000008$ | $0.0018 \pm 0.0002$ | 0.09 |
| $K_i = k_d/k_a$ | nM | $8.4 \pm 0.2$ | $3.8 \pm 0.2$ | 2.2 |
| $t_{\text{res}}$ | min | $104 \pm 5$ | $9 \pm 1$ | 11.5 |

The results of a comprehensive kinetic analysis using multiple replicates (*n* = 3, for each inhibitor) are summarized in **Table 1** (see **Table S3-S4** for detailed analysis), where *k*_a_ is the association rate constant and *k*_d_ is the dissociation rate constant. The inhibition constant *K*i and the drug-target residence time (*t*_res_) were computed from these primary regression parameters using the usual formulas (Copeland et al., 2006), while assuming that both inhibitors are kinetically competitive with the fluorogenic peptide substrate (see *Methods* for details). The results summarized in **Table 1** indicate that RAY1216 has a more than an order of magnitude (12×) lower dissociation rate constant in comparison with PF-07321332. Thus, the drug-target residence time for RAY1216 is measured in hours (1.7 hr), instead of in minutes (9 min) in the case of PF-07321332. At the same time, the equilibrium binding affinity of RAY1216 (8.4 nM) measured by the inhibition constant *K*_i_ is only approximately two-fold lower than that of PF-07321332. Note that *K*_i_ = (3.8 ± 0.2) nM reported here for PF-07321332 is in good agreement with *K*_i_ = 3.1 (1.5−6.8) nM previously reported by Pfizer (Owen *et al*., 2021). The observed enzyme inhibition kinetics, in particular the drug-target residence time results listed in **Table 1**, is consistent with slow-tight inhibition of M^pro^ by RAY1216, suggesting that RAY1216 forms a more stable enzyme-inhibitor complex (E-I) than that formed by PF-07321332.

## Structure of RAY1216 bound to SARS-CoV-2 M^pro^

To further understand the activity of RAY1216, we soaked SARS-CoV-2 M^pro^ crystals with 6 mM RAY1216 in crystallization solution and the structure of RAY1216 bound to M^pro^ at 2.0 Å resolution was determined by X-ray diffraction **(Fig. 3a** and **Table S5)**. We identified unambiguous electron density consistent with RAY1216 molecules in both active sites of M^pro^ dimer **(Fig. 3a** and **Fig. S10)** and the dimer appears to be largely symmetric **(Fig. 3a and Fig. S11)**. The electron density shows that RAY1216 is covalently attached to M^pro^ via a thiohemiketal bond formed between the S*γ* sulfur of the catalytic Cys145 and the α-keto carbon of the RAY1216 warhead **(Fig. 3b** and **Fig. S10)**. The α-ketoamide warhead at the inhibitor P1’ position is able to interact with the M^pro^ active site through a number of potential hydrogen bonds: the oxyanion (or hydroxyl) group of the thiohemiketal accepts a hydrogen bond from His41; and the warhead amide oxygen is within hydrogen bond accepting distance of the backbone amides of Gly143, Ser144 and Cys145 which form the canonical cysteine protease “oxyanion hole” **(Fig. 3c)**. These interactions are consistent with the proposal that the *α*-ketoamide represents a superior warhead through its ability to engage two hydrogen bonding interactions to the target protease catalytic center, rather than just one (Zhang *et al*., 2020), as seen for aldehyde (Zhang *et al*., 2020; Zhu et al., 2011) or Michael acceptor (Tan et al., 2013; Zhang *et al*., 2020) warheads. The cyclopentyl substituent on the warhead amide is well defined by the electron density (**Fig. 3b** and **Fig. S10**) and is situated 4.2 Å from the sidechain of M^pro^ Leu27, demonstrating a hydrophobic contact between the cyclopentyl moiety and the aliphatic Leu27 sidechain (**Fig. 3c**).

**Fig. 3.**
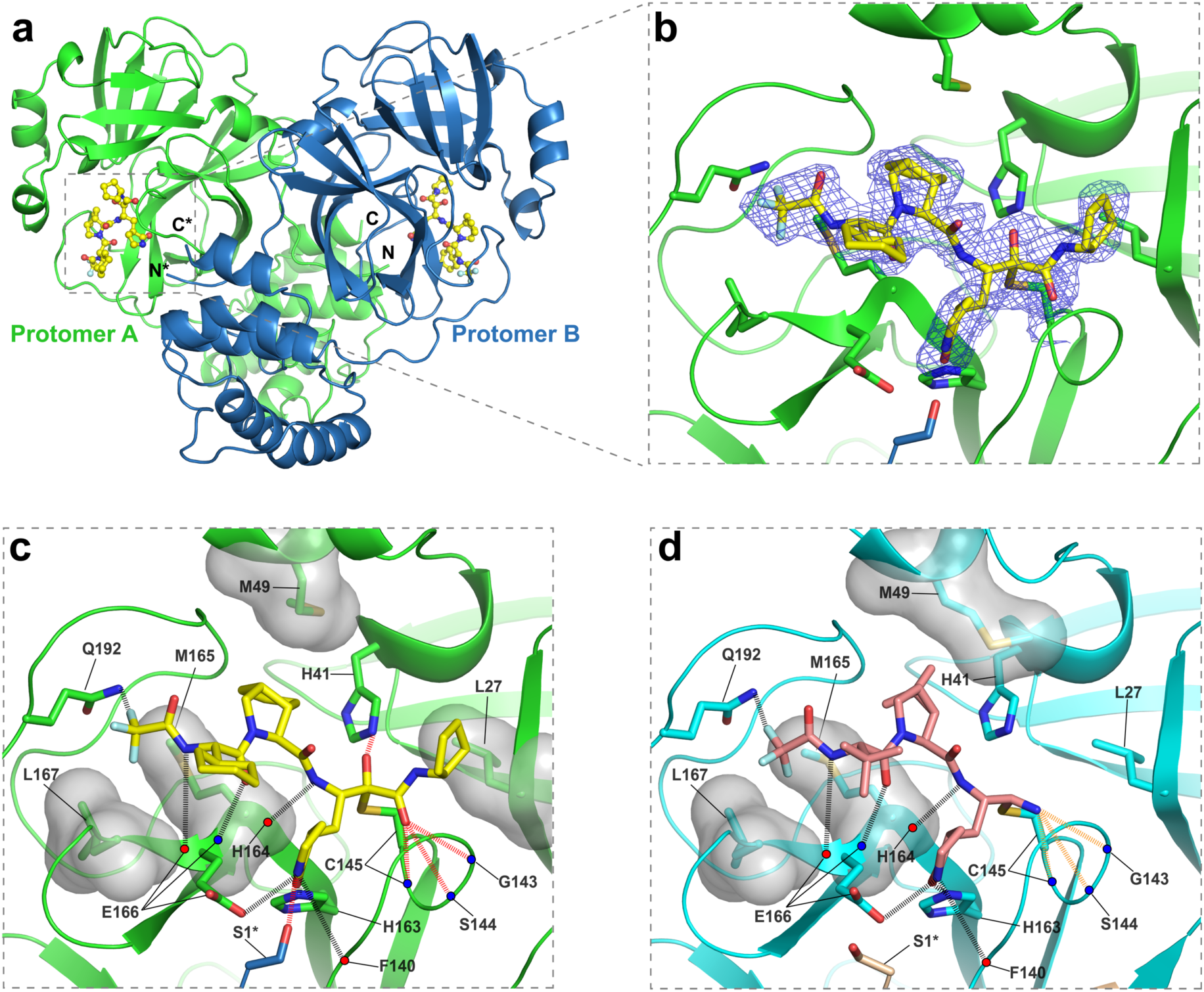
Crystal structure of RAY1216 in complex with SARS-CoV-2 M^pro^. **a**, Cartoon representation of the dimeric M^pro^ bound to RAY1216. Promoter A is in green, protomer B is in blue and RAY1216 is shown as yellow ball-and-stick models in active sites of both M^pro^ protomers. **b**, A zoom-in view of the RAY1216 bound active site of protomer A. 2Fo-Fc density map (blue mesh, contoured at 1.3σ) is shown around bound RAY1216 and the catalytic Cys145 side chain (also see **Fig. S10** for omit map densities). Clear electron density is observed for the thiohemiketal bond formed between the bound RAY1216 α-keto carbon and the catalytic Cys145 sulfur. **c,** Same view as in **b** showing detailed interactions between RAY1216 and active site of M^pro^. Selected sidechains of interacting residues are shown, backbone carbonyl and amide are represented as red and blue dots. **d**, Detailed interactions between PF-07321332 and active site of M^pro^ (based on PDB: 7RFW (Owen *et al*., 2021)) are shown in the same view as in **c**. In **c** and **d**, Molecular surfaces of selected residues involved in hydrophobic contacts with bound inhibitors are shown. Hydrogen bonds are shown in dashed lines. Extra hydrogen bonds formed by RAY1216 to M^pro^ or hydrogen bonds of different properties to M^pro^ between RAY1216 and PF-07321332 are highlighted with colours.

In the P2 position of RAY1216, the peptide bond is stabilized within the cyclopentylproline moiety previously utilized at the P2 position of telaprevir (Lin et al., 2006; Qiao *et al*., 2021). Electron density shows that the hydrophobic cyclopentyl ring slots snugly into the groove between M49 and M165 (**Fig. 3b-c** and **Fig. S10**). Plasticity has been observed for the S2 substrate binding pocket which accommodates the P2 moiety upon inhibitor binding (**Fig. S11**) (Kneller et al., 2020). It has been shown that S2 pockets in coronavirus M^pro^ have a strong preference towards hydrophobic amino acids, particularly leucine (Rut et al., 2021; Sacco et al., 2020; Xiong *et al*., 2021). It has also been shown in a separate study that dimethylcyclopropylproline and cyclopentylproline, used in boceprevir and telaprevir respectively (**Fig. 1**), when incorporated in *α*-ketoamide inhibitors, can each occupy the S2 pocket with similar potencies (Qiao *et al*., 2021).

The P3 moiety of RAY1216 features a cyclohexyl group that extends towards the exterior of the active site without making any direct contacts with M^pro^ (**Fig. 3c**). The density for the cyclohexyl *para*-carbon positioned furthest from the active site cavity is weak (**Fig. 3b**), suggesting that the cyclohexyl group remains relatively flexible within the inhibitor-enzyme complex. Nevertheless, it has been reported that substituents at P3 position can affect both drug potency and pharmacokinetic properties (Owen *et al*., 2021; Qiao *et al*., 2021).

RAY1216 and PF-07321332 share the same γ-lactam and tri-fluoroacetamide moieties at P1 and P4 respectively. The P1 γ-lactam is known as an optimal fragment for viral protease inhibition as it mimics glutamine and has been proven to be responsible for potent inhibitory activity against a variety of enzymes with specificity towards native substrates with a P1 glutamine (Dragovich et al., 1999; Owen *et al*., 2021; Qiao *et al*., 2021). In the RAY1216: M^pro^ complex, the γ-lactam nitrogen donates potential hydrogen bonds to the backbone carbonyl oxygen of Phe140 (3.19 Å), to the carboxylate of Glu166 (3.17 Å), and to the sidechain hydroxyl of Ser1 from the second monomer of the M^Pro^ dimer (**Fig. 3c**). The γ-lactam carbonyl oxygen accepts a hydrogen bond (2.54 Å) from the imidazole of His163 (**Fig. 3c**). These interactions have also been observed in the complex formed between PF-07321332 and M^pro^ (**Fig. 3d**) (Owen *et al*., 2021). Clear electron density is observed for the P4 tri-fluoroacetamide capping moiety in the RAY1216:M^pro^ complex structure (**Fig. 3b** and **Fig.S10**), it contacts Leu167 sidechain and accepts a hydrogen bond from Gln192 amide (**Fig. 3c**). Equivalent interactions have been observed in the PF-07321332:M^pro^ complex structure (**Fig. 3d**) (Owen *et al*., 2021). In summary, despite differences in the P1’ warhead, P2 bicycloproline and P3 substituent structures, interactions mediated by the P1 γ-lactam and P4 tri-fluoroacetamide moieties are largely maintained between RAY1216 and PF-07321332.

## Antiviral activities of RAY1216 in cell culture and mouse models

Based on the encouraging *in vitro* activity of RAY1216, we next sought to investigate inhibitory activity of RAY1216 towards SARS-CoV-2 infection in cell and mouse model. The 50% cytotoxic concentration (CC_50_) of RAY1216 was determined to be 511 μM for VeroE6 cells (**Fig. S12**). In virus inhibition assays the half-maximal effective concentration (EC_50_) values for RAY1216 against different SARS-CoV-2 variants are 95 nM (WT), 130 nM (Alpha), 277 nM (Beta), 97 nM (Delta), 86 nM (Omicron BA.1) and 158 nM (Omicron BA.5), respectively (**Fig. 4a**). The corresponding selectivity indices (SI, CC_50_/EC_50_) are ∼5380 (WT), ∼3930 (Alpha), ∼1850 (Beta), ∼5270 (Delta), ∼5940 (Omicron BA.1) and ∼3230 (Omicron BA.5), respectively **(Table 2 and Fig. S13)**.

**Fig 4.**
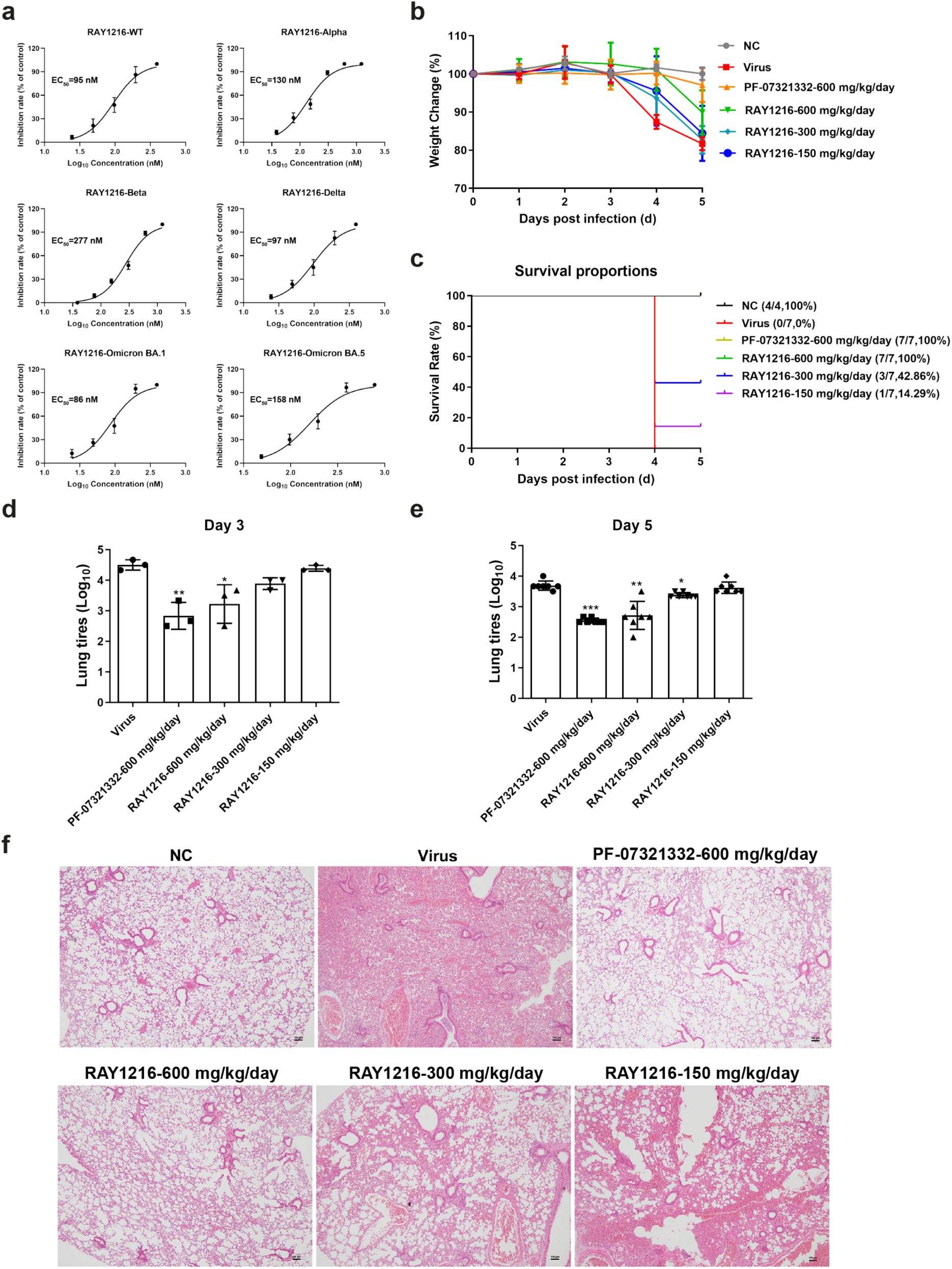
Antiviral activities of RAY1216 in cell culture and animal model. **a**, Inhibition of SARS-CoV-2 wildtype ancestral strain and variants in cell culture. Protection of Vero E6 cell from cytopathic effect (CPE) of SARS-CoV-2 virus infection was assessed by MTT cell viability assay (mean ± SD, *n* = 3). Virus inhibition titres are estimated from dose response curves of cell survival vs RAY1216 concentration. **b** and **c**, body weight change (mean ± SD) and survival rates of ACE2 transgenic C57BL/6 mice infected with SARS-CoV-2 after receiving indicated daily oral doses of RAY1216, PF-07321332 or PBS control (*n* = 7). **d** and **e**, SARS-CoV-2 virus titres (mean ± SD) in mouse lung tissues at 3 d.p.i. (*n* = 3) and 5 d.p.i (*n* = 7) after receiving indicated daily doses of RAY1216 or PF-07321332. **f**, Comparison of virus induced histology changes in mouse lung tissues after receiving indicated oral daily doses of RAY1216, PF-07321332 (*n* = 3). Histology examples of no virus (NC) and virus (virus) controls are included for comparison.

**Table 2.** Antiviral activities of RAY1216 compared to PF-07321332 in cell culture.

| SARS-CoV-2<br>variant | RAY1216 (nM) |  | PF-07321332 (nM) |  |
| --- | --- | --- | --- | --- |
|  | EC<sub>50</sub> | EC<sub>90</sub> | EC<sub>50</sub> | EC<sub>90</sub> |
| WT | 95 | 228 | 94 | 193 |
| Alpha | 130 | 349 | 72 | 205 |
| Beta | 277 | 685 | 145 | 328 |
| Delta | 97 | 251 | 71 | 177 |
| Omicron BA.1 | 86 | 204 | 107 | 215 |
| Omicron BA.5 | 158 | 359 | 111 | 266 |

We further characterized the protective effect of RAY1216 against virus infection in a human ACE2 transgenic mouse model (Bao et al., 2020). Mice were intranasally challenged with lethal doses (10^5^ PFU) of SARS-CoV-2 (Delta variant) and the protective effect of RAY1216 was assessed. The mortality of the mice in the untreated virus-infected group was 100% at 5 days post-infection. RAY1216 administered at three different doses (600 mg/kg/day, 300 mg/kg/day and 150 mg/kg/day) was able to protect mice infected with SARS-CoV-2 by 100%, 43% and 14%, respectively (**Fig. 4b**). This result suggests that treatment with RAY1216 effectively prolonged survival of SARS-CoV-2 infected mice (**Fig. 4c**). To examine effect of RAY1216 on lung virus titre and pathology, a separate set of experiments was performed with a non-lethal dose of virus inoculum (10^3.5^ PFU). RAY1216 (600 mg/kg/day and 300 mg/kg/day) decreased viral titres in lungs significantly compared with the infection-only group (**Fig. 4d**). Compared to the infection-only group, RAY1216 (600 mg/kg/day) was able to reduce lung virus titre by more than 1 Log unit. This effect may be slightly weaker for RAY1216 compared to PF-07321332 under the same experimental set-up (**Fig. 4d and e**), but the difference is not statistically significant. Lung histopathology of infected mice, compared to that of infected mice treated by RAY1216, shows that RAY1216 administered at 600 mg/kg/day and 300 mg/kg/day reduced virus induced pathology (**Fig. 4f**). RAY1216 administered at a dose of 600 mg/kg/day provided a similar level of protection against lung tissue inflammation injury to that observed with PF-07321332 (**Fig. 4f**).

## Pharmacokinetics of RAY1216

Pharmacokinetics (PK) can significantly influence drug therapeutic efficacy. We next examined pharmacokinetics of RAY1216 and PF-07321332 in head-to-head experiments. RAY1216 and PF-07321332 show comparable *in-vitro* stabilities in plasmata of various different animal species (**Table S6**), based on this result, we investigated *in-vivo* PK properties of RAY1216 and compared with those of PF-07321332 in mice, rats and cynomolgus macaques **(Table 3)**. Following intravenous (IV) administration, RAY1216 has plasma clearance (Cl) rates in the range of 10 – 22.5 mL/min/kg (compared to 23.4 – 30.2 mL/min/kg for PF-07321332) and elimination half-lives in the range of 0.9 - 3.8 h (compared to 0.3 – 0.7 h for PF-07321332) among different animals. Following oral (PO) administration, RAY1216 has elimination half-lives ranging between 2.6 – 14.9 h (compared to 1.1-1.4 h for PF-07321332) among the different animal models. These characteristics represent an improvement over PF-07321332, which demonstrates faster plasma clearance and shorter elimination half-lives under equivalent conditions across all the animal models tested. These *in vitro* and *in vivo* data indicate that RAY1216 may have promising human PK profile. Indeed, it has been noted in a number of studies that α-ketoamides appear to possess superior chemical and metabolic stability, particularly comparing to aldehyde based peptidomimetics (Robello et al., 2021).

**Table 3.** Compound pharmacokinetics parameters in different animal species. **C_max_:** the maximum observed concentration of the drug collected in bodily material from subjects in a clinical study **T_max_:** the time it takes to reach the maximum concentration or time to C_max_ **AUC:** “Area Under the Curve” and represents the total exposure of the drug experienced by the subject in a clinical study **Cl:** total plasma clearance **Vd_ss_:** Steady state volume of distribution **T_1/2_:** Half-time is the time it takes for half the drug concentration to be eliminated **oral (F%):** Oral bioavailability

| Compound | Species | dose<br>(mg/kg) | C<sub>max</sub><br>(nM) | T<sub>max</sub><br>(h) | AUC<sub>0~last</sub><br>(nM·h) | Cl<br>(mL/min/kg) | Vd<sub>ss</sub><br>(L/kg) | T<sub>1/2</sub><br>(h) | oral F<br>(%) |
| --- | --- | --- | --- | --- | --- | --- | --- | --- | --- |
| RAY1216 | mouse | 3.0 (IV) | -- | -- | 7789 | 10 | 1.4 | 3.8 | -- |
|  |  | 10 (PO) | 1287 | 2.0 | 5698 | -- | -- | 2.6 | 22 |
|  | rat | 2.0 (IV) | -- | -- | 4505 | 12.5 | 1.1 | 2.2 | -- |
|  |  | 10 (PO) | 916 | 0.9 | 7429 | -- | -- | 4.3 | 33 |
|  | cynomolgus<br>macaque | 1.0 (IV) | -- | -- | 1157 | 22.5 | 1.0 | 0.9 | -- |
|  |  | 5.0 (PO) | 102 | 1.5 | 458 | -- | -- | 14.9 | 8 |
| PF-07321332 | mouse | 3.0 (IV) | -- | -- | 3327 | 30.2 | 0.46 | 0.7 | -- |
|  |  | 10 (PO) | 2715 | 0.3 | 3577 | -- | -- | 1.1 | 33 |
|  | rat | 2.0 (IV) | -- | -- | 1600 | 42 | 1.2 | 0.8 | -- |
|  |  | 10 (PO) | 2173 | 0.4 | 2384 | -- | -- | 1.0 | 30 |
|  | cynomolgus<br>macaque | 1.0 (IV) | -- | -- | 1511 | 23.4 | 0.5 | 0.3 | -- |
|  |  | 5.0 (PO) | 601 | 1.5 | 793 | -- | -- | 1.4 | 11 |

In mouse pharmacokinetics experiments, RAY1216 also exhibits improvements when compared to PF-07321332, in area under the curve from time 0 extrapolated to last (AUC_0∼last_) for serum drug concentration (**Fig. 5**). Based on EC50/EC90 values determined using VeroE6 cells, a single IV dose of 3 mg kg^−1^ maintained the RAY1216 plasma concentration above EC50 and EC90 for 4 hours and 8 hours respectively. A single PO dose of 10 mg kg^−1^ maintained the RAY1216 plasma concentration above EC50 and EC90 for 6 hours and 10 hours respectively. Both parameters represent a marked improvement over those obtained for PF-07321332, the active anti-viral component in Paxlovid.

**Fig 5.**
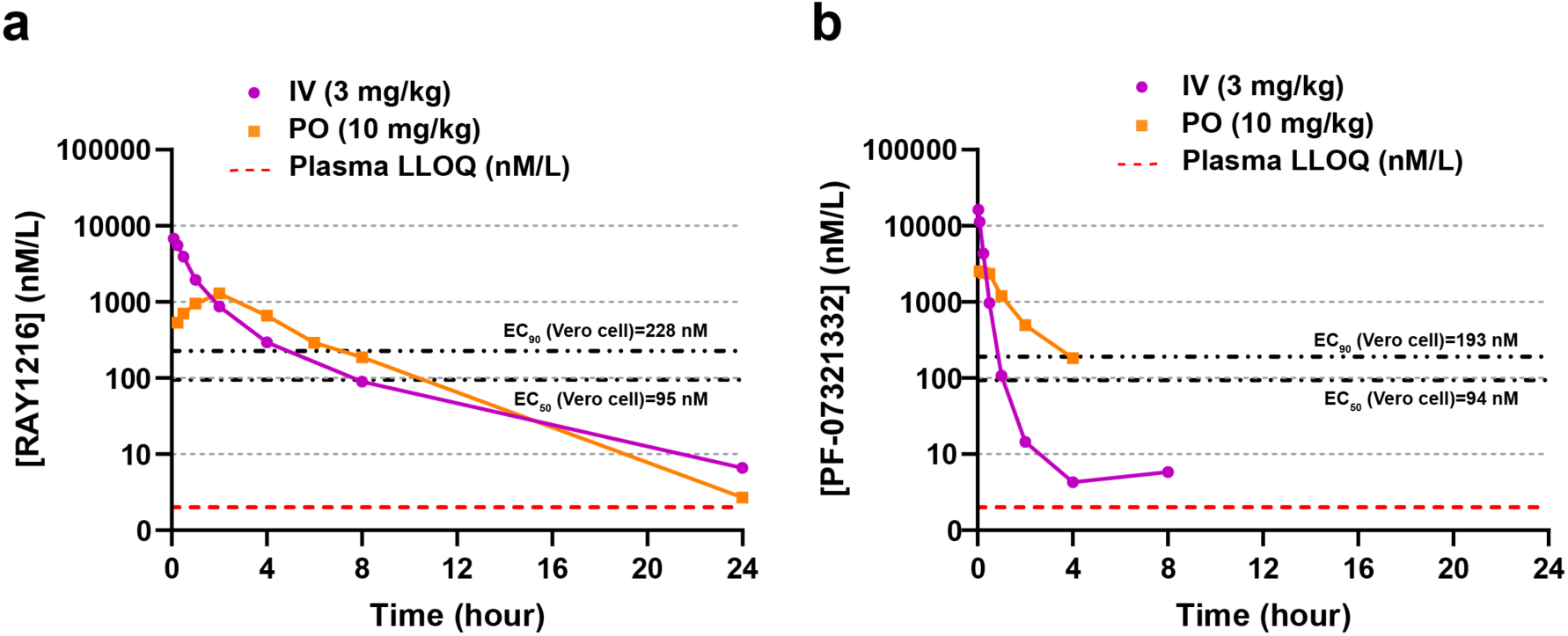
Plasma concentrations of RAY1216 and PF-07321332 after intravenous injection (IV) dosing and gavage (PO) dosing in mice. Red dashed line represents the lower limit of quantitation (LLOQ: 2 nM/L) of plasma drug concentration. EC_50_ and EC_90_ values against SARS-CoV-2 WT strain determined using Vero E6 cell are indicated by black dashed lines.

## Discussion

In this study, we characterized inhibition of SARS-CoV-2 M^pro^ by RAY1216, a peptidomimetic inhibitor. This compound features a cyclopentyl-substituted α-ketoamide warhead, a P1 pyroglutamine (known to confer selectivity towards CoV M^pro^), a P2 cyclopentylproline as originally utilized in the anti-HCV drug telaprevir (Yip *et al*., 2004a), a P3 cyclohexylglycine, and finally a P4 tri-fluoroacetamide as utilized in PF-07321332. We found more extensive interactions between the covalently attached drug and the SARS-CoV-2 M^pro^ enzyme in the crystal structure of the RAY1216:M^pro^ acyl enzyme complex, compared with that of PF-07321332. In enzyme inhibition assays we found that RAY1216 inhibits M^pro^ via a slow-tight mechanism, with an approximately 12-fold longer drug-target residence time. These inhibition characteristics suggest that RAY1216 forms a more stable acyl-enzyme adduct when compared with PF-07321332. It has recently emerged that drug-target residence time is an important parameter to optimise for drug efficacy (Copeland *et al*., 2006; Dahl and Akerud, 2013; Lu and Tonge, 2010). In pharmacokinetic studies, RAY1216 showed improved elimination half-lives compared to PF-07321332. This may allow its use without ritonavir which is known to have significant unwanted drug-drug interactions.

In summary, RAY1216 possesses superior drug-target residence time and pharmacokinetic properties when compared with PF-07321332 (nirmatrelvir), the active anti-viral component in Paxlovid. On the other hand, PF-07321332 is slightly favoured over RAY1216 in reducing mouse lung viral titre. The real-world efficacy of RAY1216 as a potential therapeutic for the treatment of COVID-19 in humans will be revealed by phase III clinical trial, which is currently ongoing.

## Author contributions

Z.Y., X.X. and X.C. conceived the study under the direction of N.Z.; X.C., J.H., H.L., C.L. and S-H. C. provided the M^pro^ inhibitors, collected chemical characterization data and *in-vivo* and *in-vitro* pharmacokinetics data. X.H. expressed, purified M^pro^ and performed enzyme kinetics assays under the supervision of X.X.; Q.M. performed virus inhibition assays in cell culture and animal models and prepared figures with assistance from B.L., H.J., W.Z., C.Y., S.W. and under the supervision of Z.Y.; P.K. and X.X. analysed enzyme kinetics data and prepared figures; X.H. and B.Z. obtained M^pro^ crystals and performed crystal soaking experiments under the supervision of X.X.; J.X., X.H., B.Z., Y.S., Y.G. performed M^pro^ crystal diffraction experiments and collected diffraction data under the supervision of Y.X. and J.L.; X.H determined the M^pro^ crystal structures and built molecular models with assistance from C.N., L.X. and under the supervision of X.X.; H.X. and X.X. analysed the M^pro^ crystal structures and prepared figures; with input from all authors, X.X., P.K., X.H., Q.M. and X.C. wrote the initial draft which was reviewed and edited by Z.Y., S-H.C., Z.X., J.S. T.Z., J.H. and W.D.; N.Z., X.X. and Z.Y. acquired funding and supervised the research.

## Competing interests

X.C., J.H., H.L. and C.L. are employees of Guangdong Raynovent Biotech Co., Ltd, which holds the patent of RAY1216.

## Acknowledgement

We thank the staffs at beamline BL19U1 of Shanghai Synchrotron Radiation Facility for assistance on data collection. This work was supported by National Multidisciplinary Innovation Team Project of Traditional Chinese Medicine (ZYYCXTD-D-202201 to Z.Y.); Guangdong Science and Technology Foundation (2022B1111060003 to Z.Y.); Guangzhou Science and Technology Planning Project (2022B01W0001 and 202102100003 to Z.Y.); Emergency Key Program of Guangzhou Laboratory (EKPG21-06 to X.X.); R&D Program of Guangzhou Laboratory (SRPG22-002 and SRPG22-003 to X.X; TL22-13 to Z.Y.). Natural Science Fund of Guangdong Province (2021A1515011289 to X.X.). X.X. acknowledges Start-up grants from the Chinese Academy of Sciences.

## Supporting Information

**Fig. S1.**
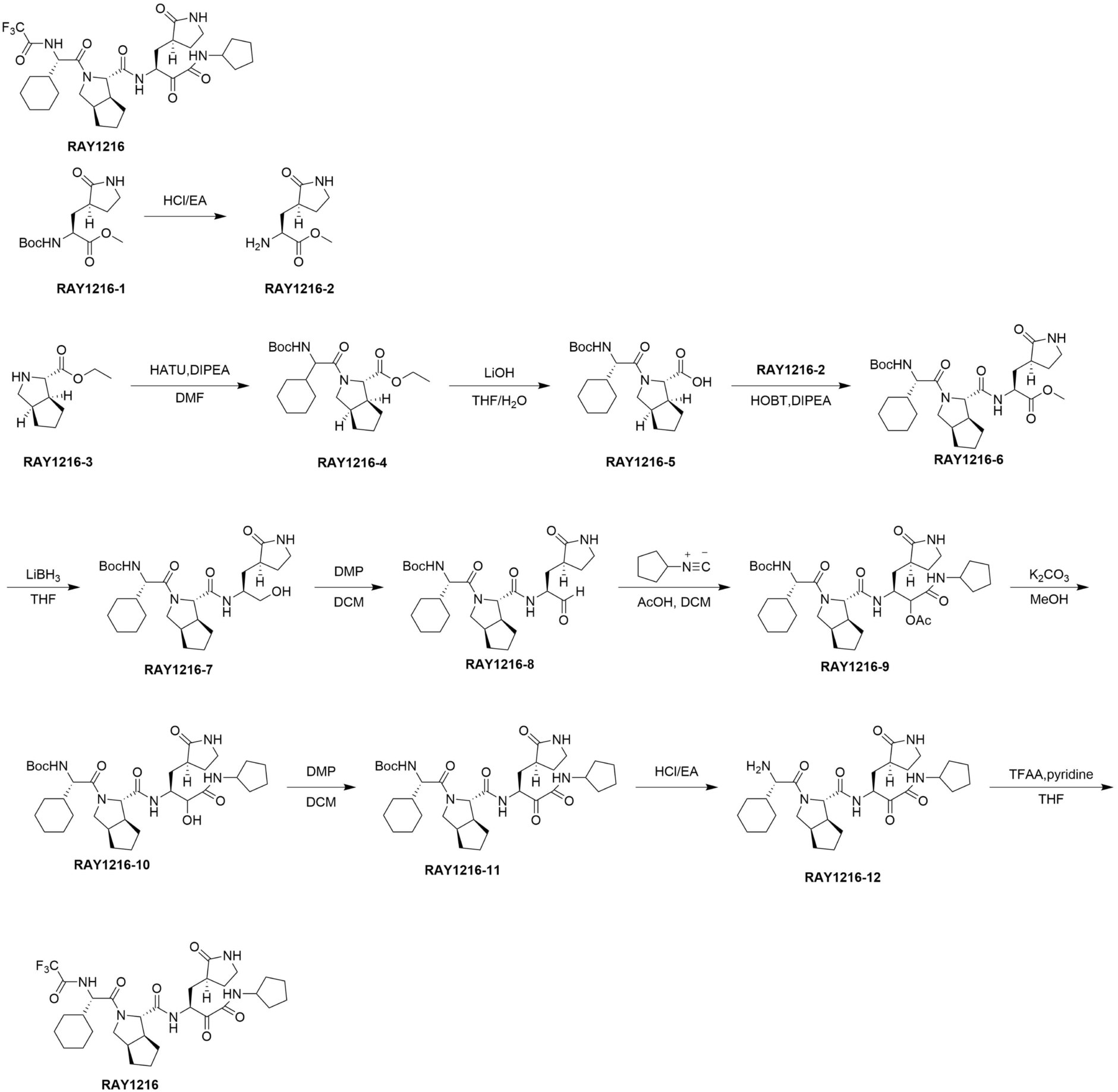
Synthesis of RAY1216.

**Fig. S2.**
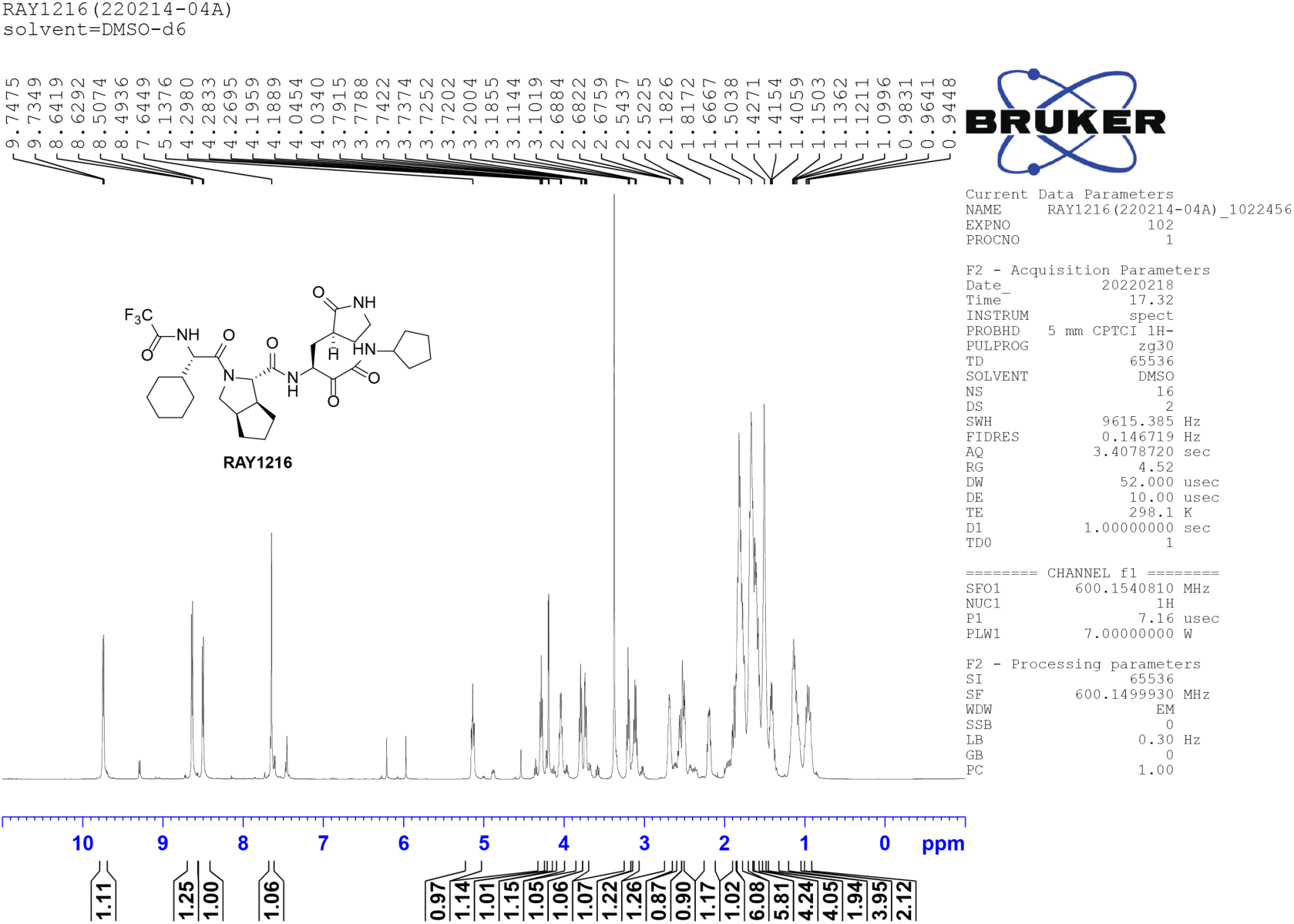
^1^H NMR spectra of RAY1216 in DMSO-d6.

**Fig. S3.**
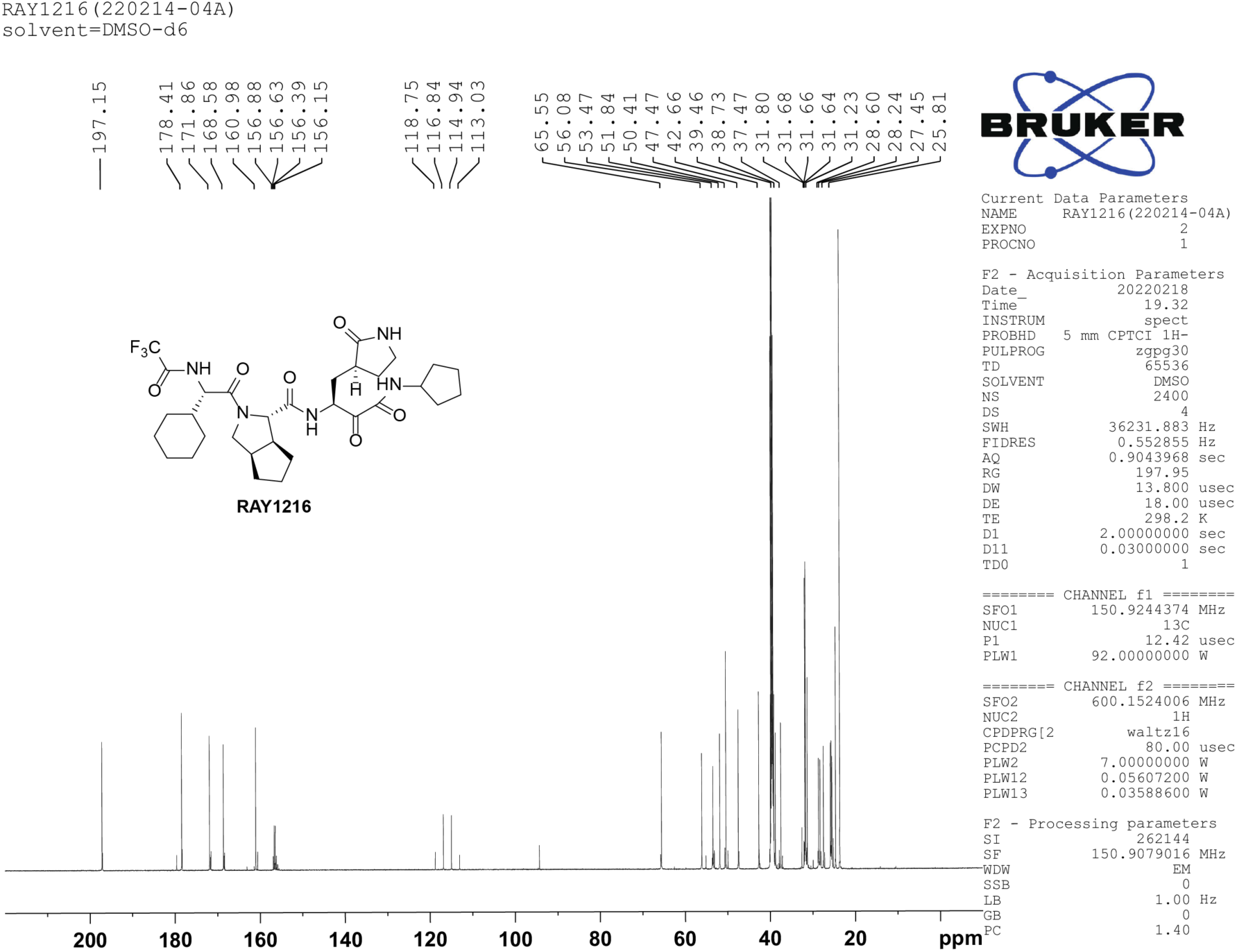
^13^C NMR spectrum of RAY1216 in DMSO-d6.

**Fig. S4.**
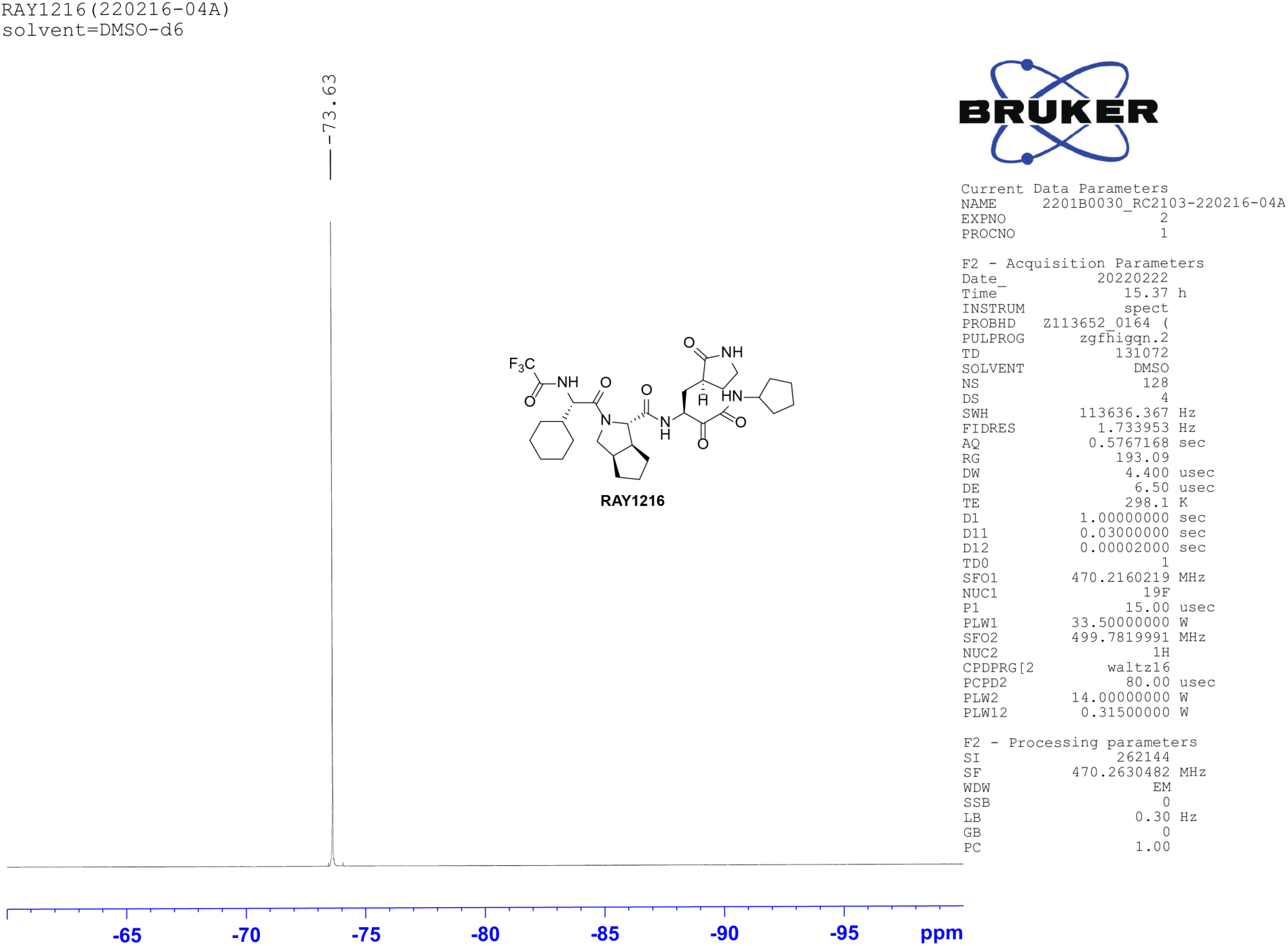
^19^F NMR spectrum of RAY1216 in DMSO-d6.

**Fig. S5.**
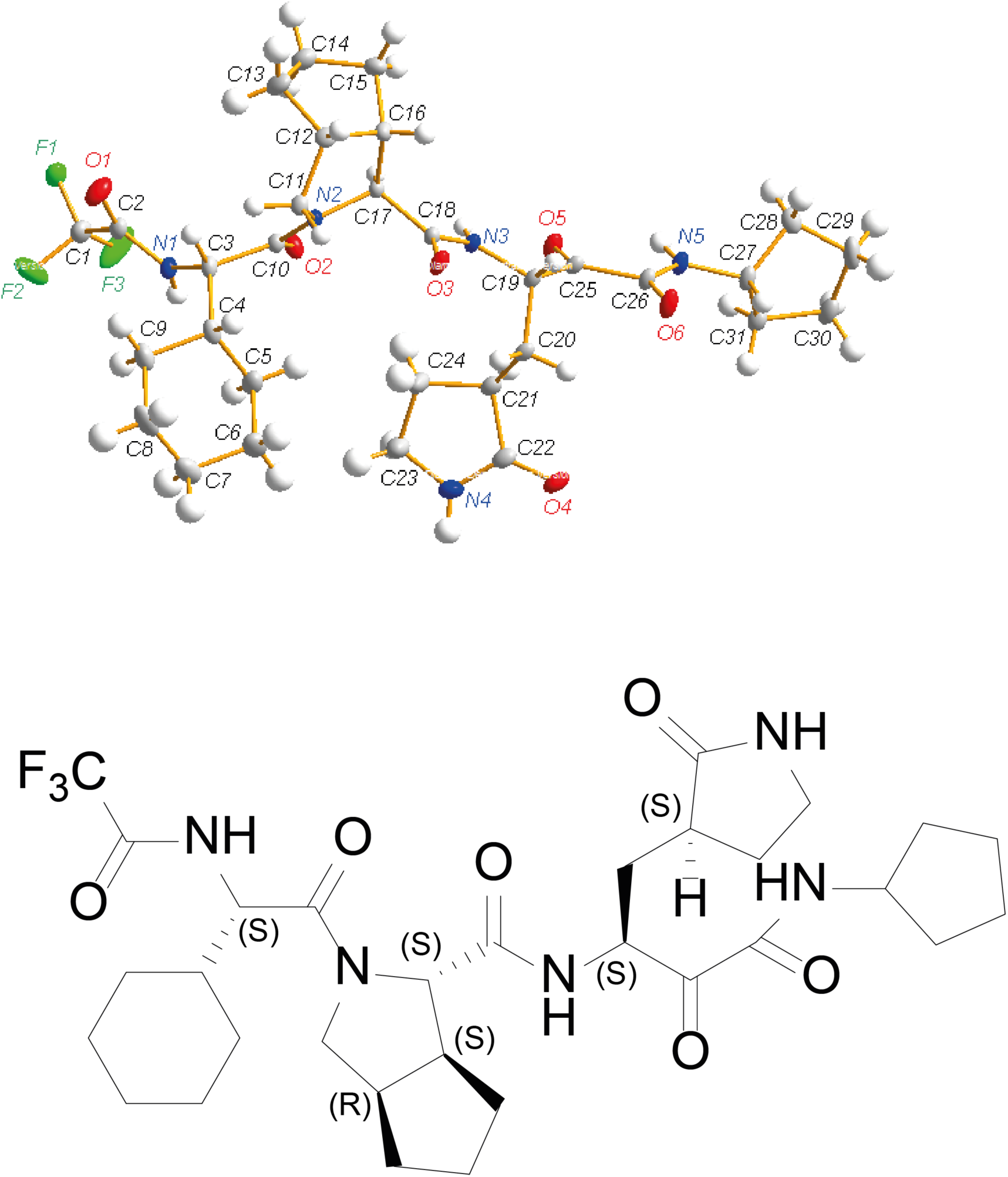
Single crystal X-ray structure of RAY1216.

**Table S1.** Data collection and statistics of single crystal X-ray structure of RAY1216.

|  |  |
| --- | --- |
| **Molecule** | RAY1216 |
| **Radiation Type** | CuKα ($\lambda$=1.54184 Å) |
| **Crystal size (mm<sup>3</sup>)** | 0.16 × 0.18 × 0.22 |
| **Crystal system** | monoclinic |
| **Space group** | *P* 2<sub>1</sub> |
| **cell dimensions** |  |
| **a, b, c (Å)** | 10.6833 9.8921 15.3665 |
| **$\alpha$, $\beta$, $\gamma$ (°)** | 90.00 91.11 90.00 |
| **Cell volume (Å<sup>3</sup>)** | 1623.63 (2) |
| **Cell formula units Z** | 2 |
| **Crystal density<sub>calc</sub> (g/cm<sup>3</sup>)** | 1.309 |
| **Crystal F (000)** | 680 |
| **Absorption coefficient ($\mu$/mm<sup>-1</sup>)** | 0.862 |
| **Index ranges** | -12 ≤ *h* ≤ 12, -11 ≤ *k* ≤ 11, -18 ≤ *l* ≤ 18 |
| **Cell measurement temperature (K)** | 149.9 (8) |
| **2θ range for data collection (°)** | 5.752 to 133.106 |
| **Goodness-of-fit on F<sup>2</sup>** | 1.041 |
| **Final R indexes [$\vert I\vert \geq 2\sigma(I)$]** | R<sub>1</sub>=0.0351, wR<sub>2</sub>=0.0914 |
| **Final R indexes [all data]** | R<sub>1</sub>=0.0358, wR<sub>2</sub>=0.0921 |
| **Largest diff. peak/hole/e Å<sup>-3</sup>** | 0.39/-0.31 |
| **Reflections collected/unique** | 40929/5638 [*R*<sub>int</sub> = 0.0391] |
| **Flack parameter** | 0.10(5) |

**Fig. S6.**
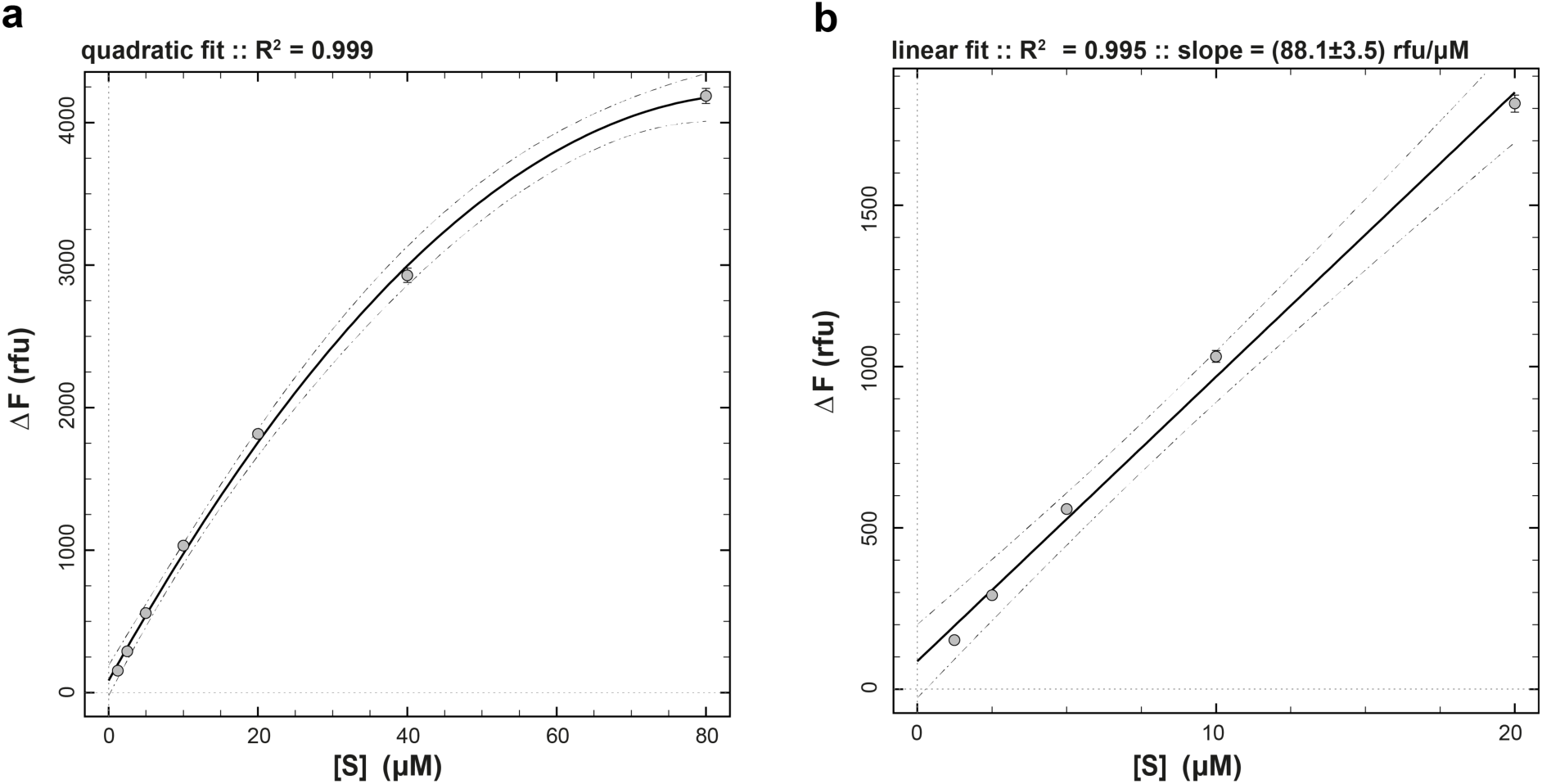
Determination of molar response coefficient of fluorescent Dabcyl-Edans cleavage product. **a**, total observed change in fluorescence intensity (Δ*F*) upon complete conversion of the substrate to product is plotted against the starting concentration of substrate ([S]) over the complete concentration range (1.25 – 80 µM); data points are fit to the quadratic function. **b**, Fit of the reduced concentration range (1.25 – 20 µM) to a straight line.

**Fig. S7.**
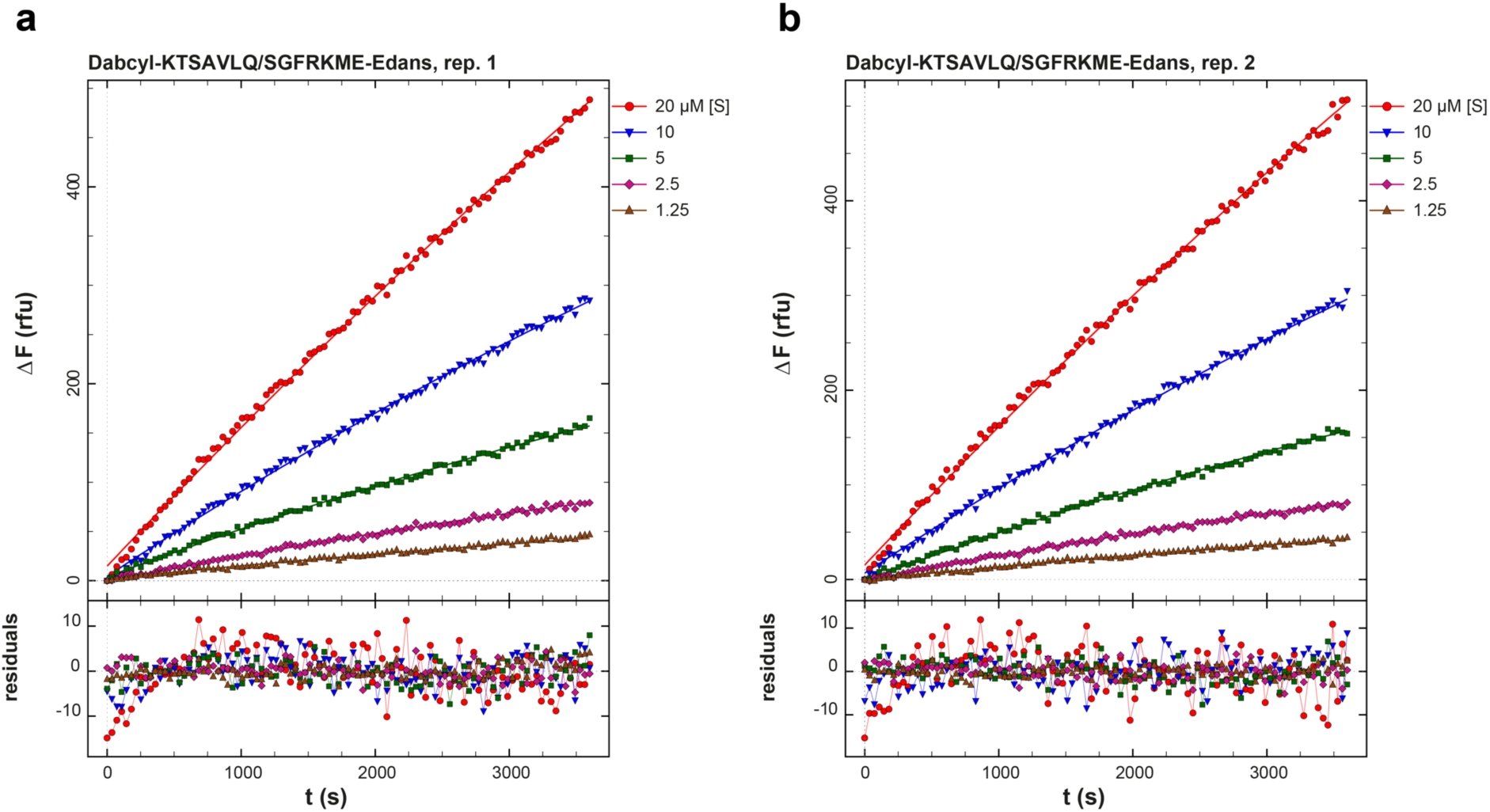
Substrate-only progress curves fit to the Michaelis-Menten reaction mechanism shown in Scheme 1. The mathematical model for the fitted curves is represented by **Eq.1 – Eq.5. a**, Replicate 1. **b**, Replicate 2.

**Table S2.**
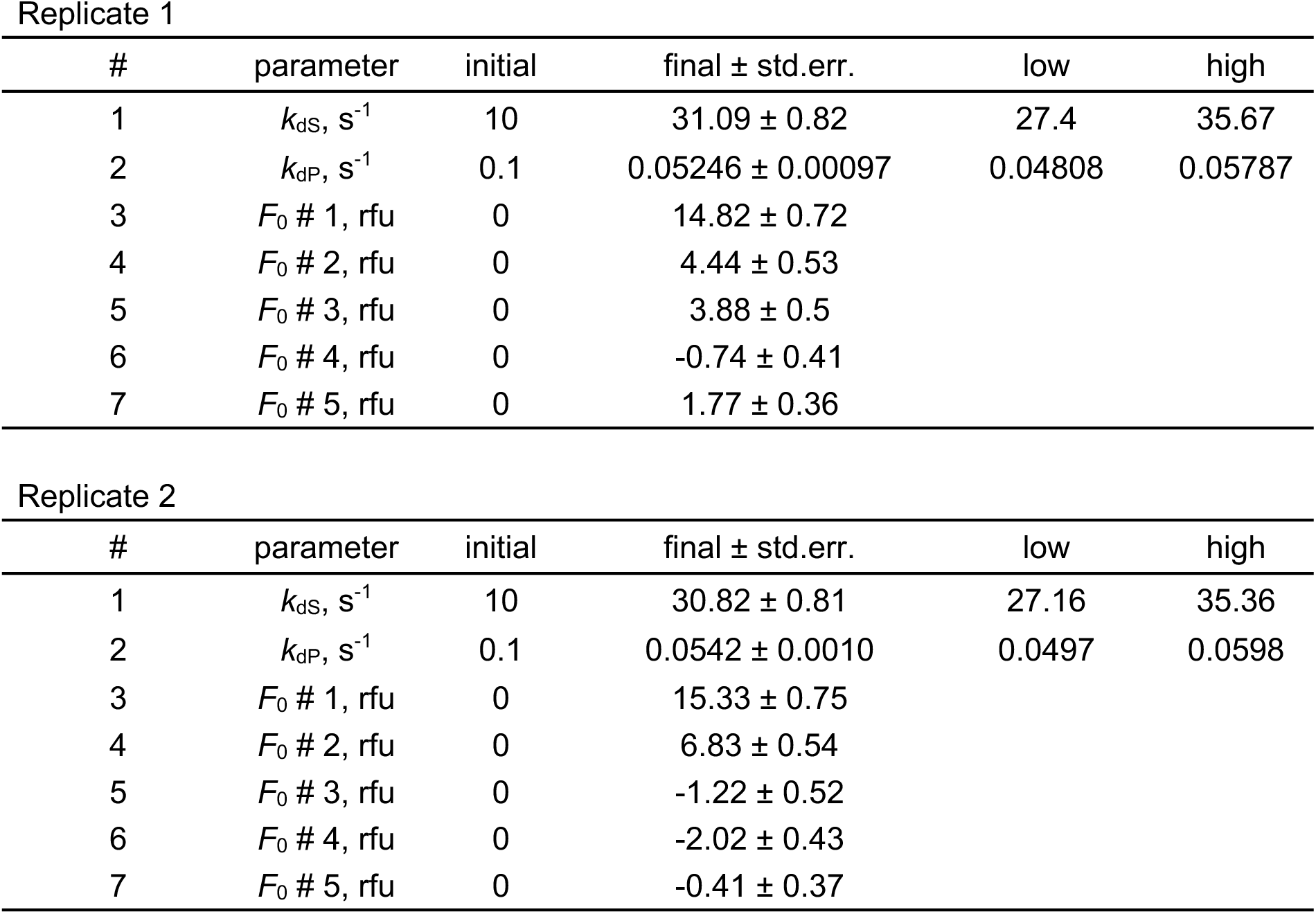
Results of fit of substrate-only progress curves to the Michaelis-Menten reaction model, replicate 1 and 2. Columns labeled as “low” and “high” contain the lower and upper limits, respectively, of non-symmetrical confidence intervals obtained by the profile-t method of Bates and Watts (Bates and Watts, 1988; Watts, 1994) while using the empirical value of ΔSSQ = 5% according to # parameter initial the previously suggested method (Johnson, 2009).

| # | parameter | initial | final ± std.err. | low | high |
| --- | --- | --- | --- | --- | --- |
| 1 | $k_{dS}, s^{-1}$ | 10 | $31.09 \pm 0.82$ | 27.4 | 35.67 |
| 2 | $k_{dP}, s^{-1}$ | 0.1 | $0.05246 \pm 0.00097$ | 0.04808 | 0.05787 |
| 3 | $F_0 \# 1, rfu$ | 0 | $14.82 \pm 0.72$ | | |
| 4 | $F_0 \# 2, rfu$ | 0 | $4.44 \pm 0.53$ | | |
| 5 | $F_0 \# 3, rfu$ | 0 | $3.88 \pm 0.5$ | | |
| 6 | $F_0 \# 4, rfu$ | 0 | $-0.74 \pm 0.41$ | | |
| 7 | $F_0 \# 5, rfu$ | 0 | $1.77 \pm 0.36$ | | |

**Table S2. Results of fit of substrate-only progress curves to the Michaelis-Menten reaction model, replicate 1 and 2.**
| # | parameter | initial | final ± std.err. | low | high |
| --- | --- | --- | --- | --- | --- |
| 1 | $k_{dS}, s^{-1}$ | 10 | $30.82 \pm 0.81$ | 27.16 | 35.36 |
| 2 | $k_{dP}, s^{-1}$ | 0.1 | $0.0542 \pm 0.0010$ | 0.0497 | 0.0598 |
| 3 | $F_0 \# 1, rfu$ | 0 | $15.33 \pm 0.75$ | | |
| 4 | $F_0 \# 2, rfu$ | 0 | $6.83 \pm 0.54$ | | |
| 5 | $F_0 \# 3, rfu$ | 0 | $-1.22 \pm 0.52$ | | |
| 6 | $F_0 \# 4, rfu$ | 0 | $-2.02 \pm 0.43$ | | |
| 7 | $F_0 \# 5, rfu$ | 0 | $-0.41 \pm 0.37$ | | |

**Fig. S8.**
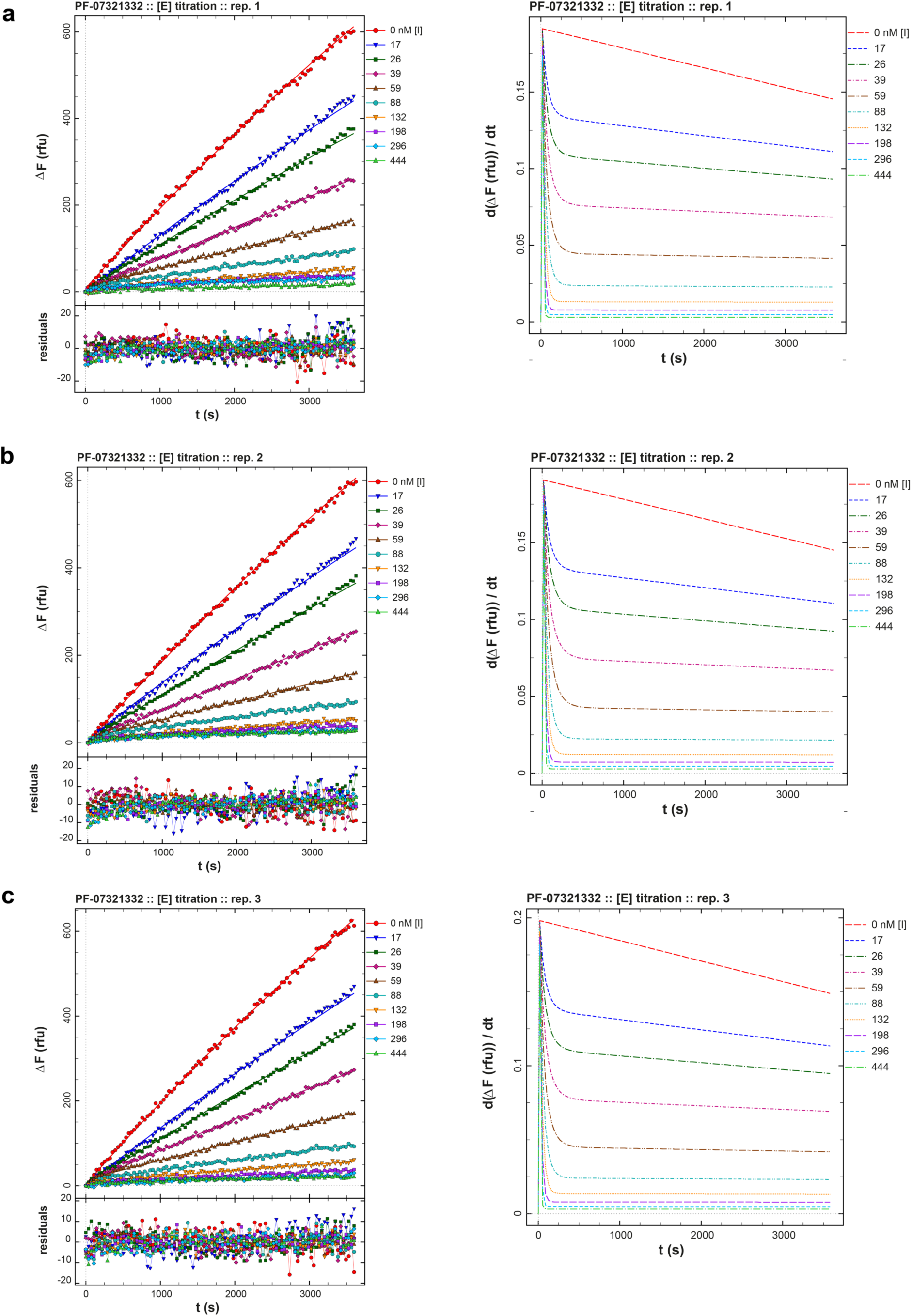
Inhibition of M^pro^ by PF-07321332. Left panels, Replicated (**a**-**c**) PF-07321332 inhibition progress curves (dots) are overlaid with best-fit model curves corresponding to the reaction mechanism in **Scheme 2** and represented by **Eq.6-Eq.11**; the residuals of the fits are shown. Right panels, corresponding plots of instantaneous reaction rates for the 3 replicates.

**Table S3.**
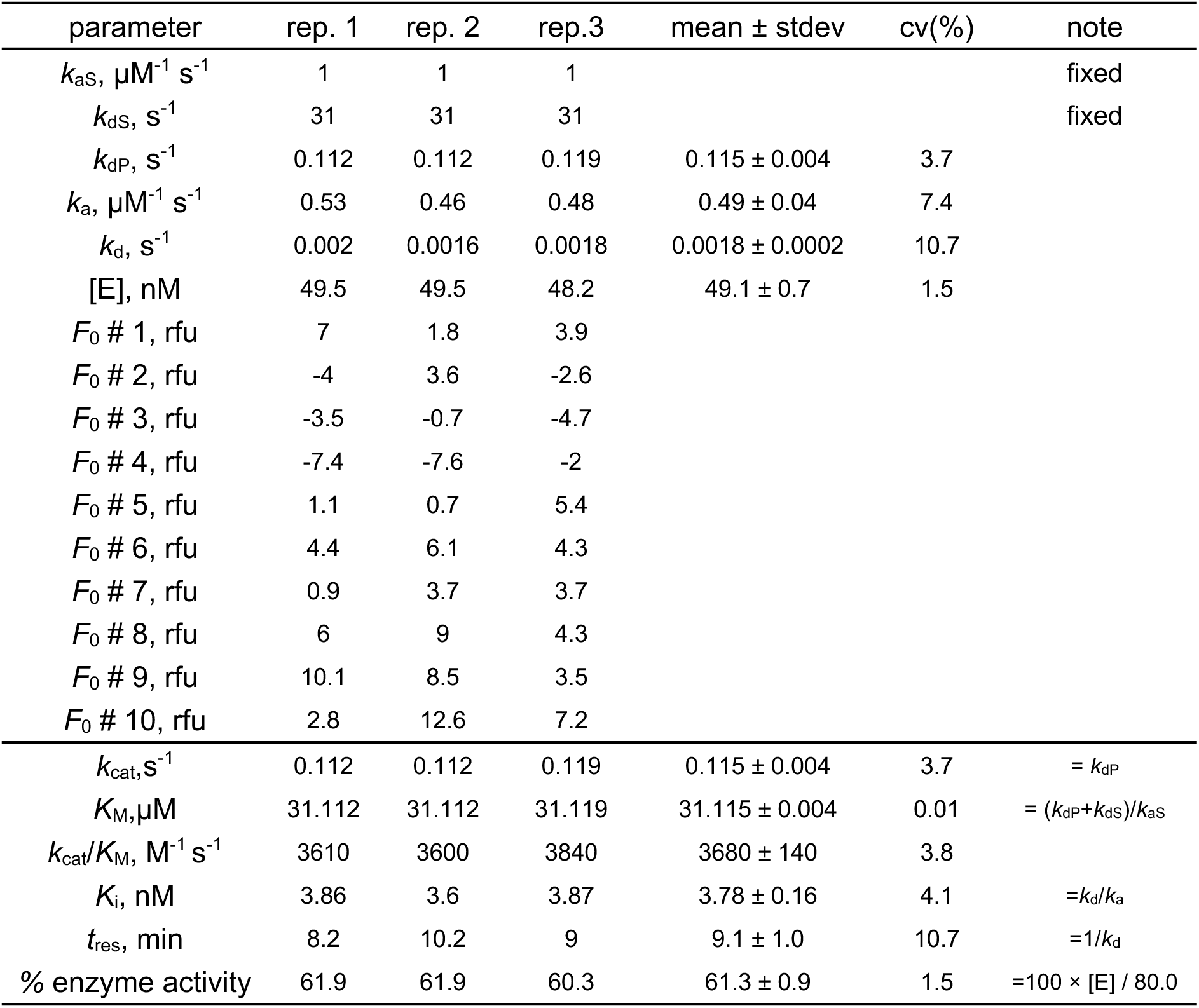
Results of fit from global analysis of PF-07321332 inhibition data. Note that both active enzyme concentration [E] (nominal concentration 80.0 nM) and the turnover number k_dP_ were treated as adjustable parameters. “stdev” is the standard deviation from replicates (n = 3) and “cv(%)” is the corresponding coefficient of variation, cv = 100 × stdev/mean. For details see Methods section.

| parameter | rep. 1 | rep. 2 | rep.3 | mean ± stdev | cv(%) | note |
| --- | --- | --- | --- | --- | --- | --- |
| $k_{aS}, \mu\text{M}^{-1} \text{s}^{-1}$ | 1 | 1 | 1 | | | fixed |
| $k_{dS}, \text{s}^{-1}$ | 31 | 31 | 31 | | | fixed |
| $k_{dP}, \text{s}^{-1}$ | 0.112 | 0.112 | 0.119 | $0.115 \pm 0.004$ | 3.7 | |
| $k_a, \mu\text{M}^{-1} \text{s}^{-1}$ | 0.53 | 0.46 | 0.48 | $0.49 \pm 0.04$ | 7.4 | |
| $k_d, \text{s}^{-1}$ | 0.002 | 0.0016 | 0.0018 | $0.0018 \pm 0.0002$ | 10.7 | |
| [E], nM | 49.5 | 49.5 | 48.2 | $49.1 \pm 0.7$ | 1.5 | |
| $F_0$ # 1, rfu | 7 | 1.8 | 3.9 | | | |
| $F_0$ # 2, rfu | -4 | 3.6 | -2.6 | | | |
| $F_0$ # 3, rfu | -3.5 | -0.7 | -4.7 | | | |
| $F_0$ # 4, rfu | -7.4 | -7.6 | -2 | | | |
| $F_0$ # 5, rfu | 1.1 | 0.7 | 5.4 | | | |
| $F_0$ # 6, rfu | 4.4 | 6.1 | 4.3 | | | |
| $F_0$ # 7, rfu | 0.9 | 3.7 | 3.7 | | | |
| $F_0$ # 8, rfu | 6 | 9 | 4.3 | | | |
| $F_0$ # 9, rfu | 10.1 | 8.5 | 3.5 | | | |
| $F_0$ # 10, rfu | 2.8 | 12.6 | 7.2 | | | |
| $k_{cat}, \text{s}^{-1}$ | 0.112 | 0.112 | 0.119 | $0.115 \pm 0.004$ | 3.7 | $= k_{dP}$ |
| $K_M, \mu\text{M}$ | 31.112 | 31.112 | 31.119 | $31.115 \pm 0.004$ | 0.01 | $= (k_{dP} + k_{dS}) / k_{aS}$ |
| $k_{cat} / K_M, \text{M}^{-1} \text{s}^{-1}$ | 3610 | 3600 | 3840 | $3680 \pm 140$ | 3.8 | |
| $K_i, \text{nM}$ | 3.86 | 3.6 | 3.87 | $3.78 \pm 0.16$ | 4.1 | $= k_d / k_a$ |
| $t_{res}, \text{min}$ | 8.2 | 10.2 | 9 | $9.1 \pm 1.0$ | 10.7 | $= 1 / k_d$ |
| % enzyme activity | 61.9 | 61.9 | 60.3 | $61.3 \pm 0.9$ | 1.5 | $= 100 \times [E] / 80.0$ |

**Fig. S9.**
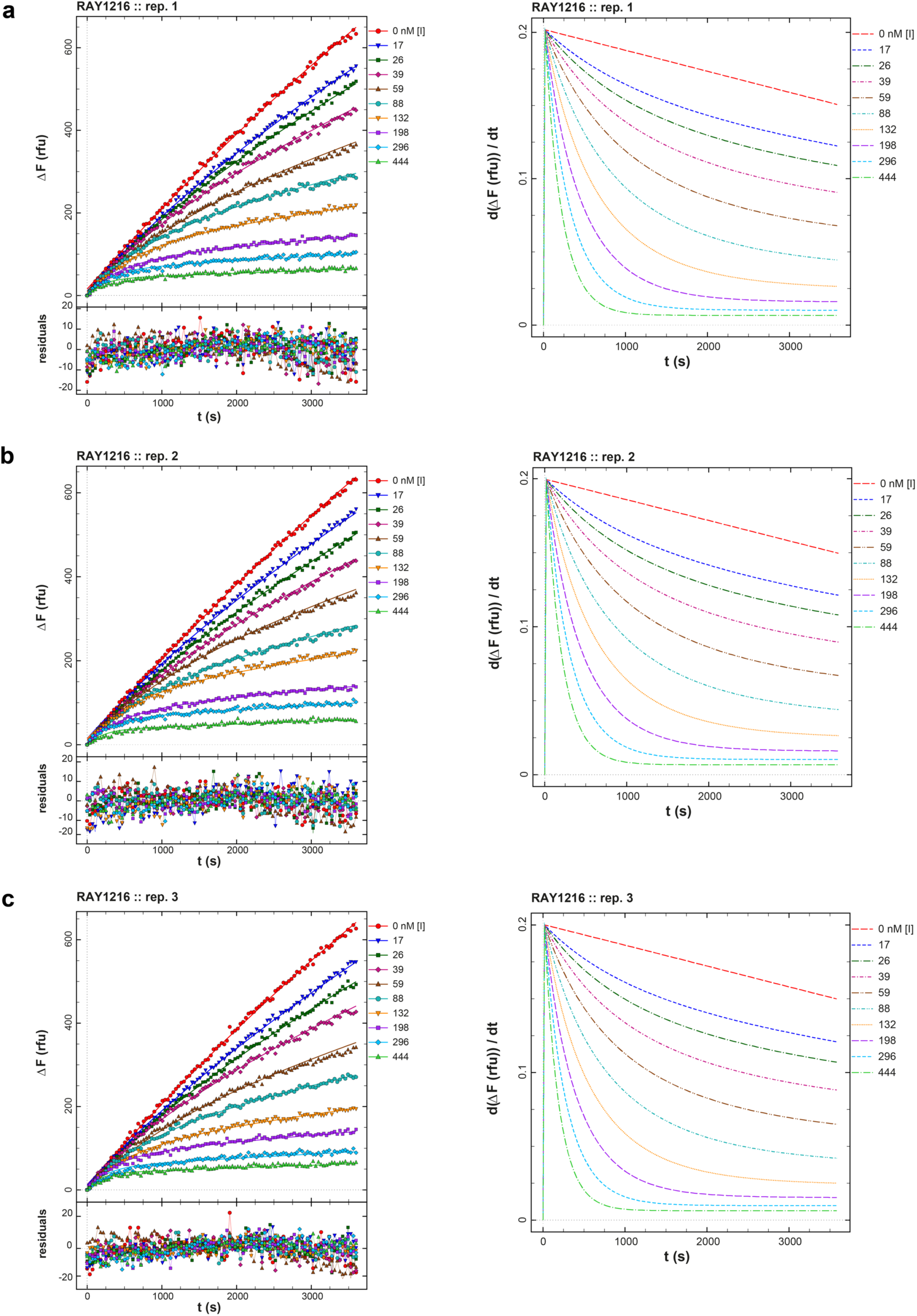
Inhibition of M^pro^ by RAY1216. Left panels, Replicated (**a**-**c**) RAY1216 inhibition progress curves (dots) are overlaid with best-fit model curves corresponding to the reaction mechanism in **Scheme 2** and represented by **Eq.6-Eq.11**; the residuals of the fits are shown. Right panels, corresponding plots of instantaneous reaction rates for the 3 replicates.

**Table S4.** Results of fit from global analysis of RAY1216 inhibition data. Note that active enzyme concentration [E] (nominal concentration 80.0 nM) was treated as adjustable parameters while the turnover number *k*_dP_ was fixed. “stdev” is the standard deviation from replicates (*n* = 3) and “cv(%)” is the corresponding coefficient of variation, cv = 100 × stdev/mean. For details see *Methods* section.

| parameter | rep. 1 | rep. 2 | rep.3 | mean ± stdev | cv(%) | note |
| --- | --- | --- | --- | --- | --- | --- |
| $k_{aS}, \mu\text{M}^{-1} \text{s}^{-1}$ | 1 | 1 | 1 | | | fixed |
| $k_{dS}, \text{s}^{-1}$ | 31 | 31 | 31 | | | fixed |
| $k_{dP}, \text{s}^{-1}$ | 0.11 | 0.11 | 0.11 | | | fixed |
| $k_a, \mu\text{M}^{-1} \text{s}^{-1}$ | 0.0183 | 0.0187 | 0.0208 | $0.0193 \pm 0.0014$ | 7.1 | |
| $k_d, \text{s}^{-1}$ | 0.000153 | 0.000161 | 0.000169 | $0.000161 \pm 0.000008$ | 4.9 | |
| [E], nM | 53.3 | 52.7 | 52.8 | $52.9 \pm 0.3$ | 0.6 | |
| $F_0$ # 1, rfu | 15.9 | 10.3 | 12.7 | | | |
| $F_0$ # 2, rfu | 11.1 | 14.7 | 12.0 | | | |
| $F_0$ # 3, rfu | 10.2 | 5.3 | 7.9 | | | |
| $F_0$ # 4, rfu | 8.5 | 2.6 | 8.7 | | | |
| $F_0$ # 5, rfu | -2.0 | 2.0 | -3.5 | | | |
| $F_0$ # 6, rfu | 5.4 | -0.9 | 3.3 | | | |
| $F_0$ # 7, rfu | 7.2 | 13.9 | 2.8 | | | |
| $F_0$ # 8, rfu | 1.1 | -1.4 | 9.4 | | | |
| $F_0$ # 9, rfu | 4.7 | 5.1 | 4.6 | | | |
| $F_0$ # 10, rfu | 0.7 | -3.2 | 5.1 | | | |
| $K_i$, nM | 8.4 | 8.6 | 8.1 | $8.4 \pm 0.2$ | 2.9 | $=k_d/k_a$ |
| $t_{\text{res, min}}$ | 109 | 104 | 99 | $104 \pm 5$ | 4.9 | $=1/k_d$ |
| % enzyme activity | 66.6 | 65.9 | 66.0 | $66.2 \pm 0.4$ | 0.6 | $=100 \times [E] / 80.0$ |

**Table S5.** Data collection and refinement statistics of M^pro^ crystal structures.

| Protein/Ligand |  | M<sup>pro</sup> free enzyme<br>(PDB: 8IGO) | M<sup>pro</sup>+RAY1216<br>(PDB: 8IGN) |
| --- | --- | --- | --- |
| Data<br>collection | wavelength (Å) | 0.97849 | 0.97853 |
|  | Space group | *C* 2 | *P* 1 |
|  | cell dimensions |  |  |
|  | a, b, c (Å) | 97.39 82.49 51.42 | 55.83 60.94 64.04 |
|  | α, β, γ (°) | 90.00 115.52 90.00 | 80.03 68.34 70.67 |
|  | Resolution (Å) | 43.46-2.00 (2.05-2.00) | 59.42-2.00 (2.05-2.00) |
|  | R<sub>merge</sub> | 0.031 (0.327) | 0.067 (0.504) |
|  | No. Reflections (total) | 156563 (11225) | 168870 (11201) |
|  | No. Reflections (unique) | 23876 (1817) | 48627 (3545) |
|  | //σ/ | 31.6 (5.1) | 9.9 (2.1) |
|  | Completeness (%) | 96.2 (98.8) | 97.5 (96) |
|  | Multiplicity | 6.6 (6.2) | 3.5 (3.2) |
|  | No. Reflections | 23851 (2089) | 47154 (4676) |
|  | R<sub>work</sub>/R<sub>free</sub> | 0.2059/0.2491 | 0.1862/0.2312 |
| Refinement | No. atoms |  |  |
|  | Protein | 2314 | 4654 |
|  | Inhibitor | - | 90 |
|  | Water | 100 | 255 |
|  | B-factors |  |  |
|  | Protein | 43.06 | 38.84 |
|  | Inhibitor | - | 37.25 |
|  | Water | 44.64 | 45.39 |
|  | R.m.s deviations |  |  |
|  | Bond lengths(Å) | 0.008 | 0.0066 |
| Validation | Bond angles(°) | 0.87 | 1.478 |
|  | MolProbity score | 1.41 | 1.94 |
|  | Clashscore | 4.15 | 6.02 |
|  | Poor rotamers (%) | 0 | 2.13 |
|  | Ramchandran plot |  |  |
|  | Favored (%) | 96.66 | 94.82 |
|  | Allowed (%) | 3.34 | 5.18 |
|  | Disallowed (%) | 0 | 0 |

**Fig S10.**
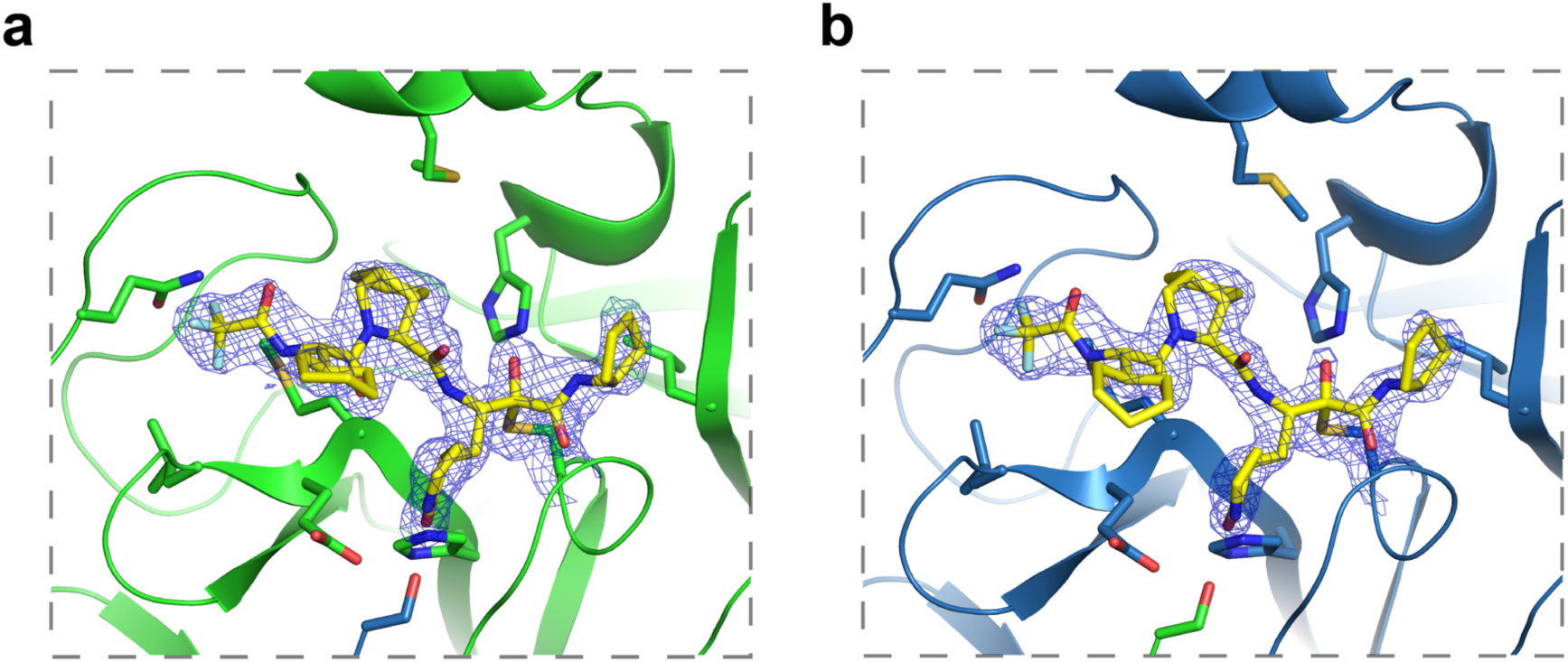
Simulated-annealing 2mFo-DFc composite omit map densities showing bound RAY1216 and covalent linkages between RAY1216 and Cys145. The simulated-annealing composite omit map for the M^pro^ dimer was calculated by omitting bound RAY1216 and Cys145 in both monomers. Shown densities are contoured at 1.1σ, models are colored according to **Fig. 2**. **a**, density in protomer A. **b**, density in protomer B.

**Fig S11.**
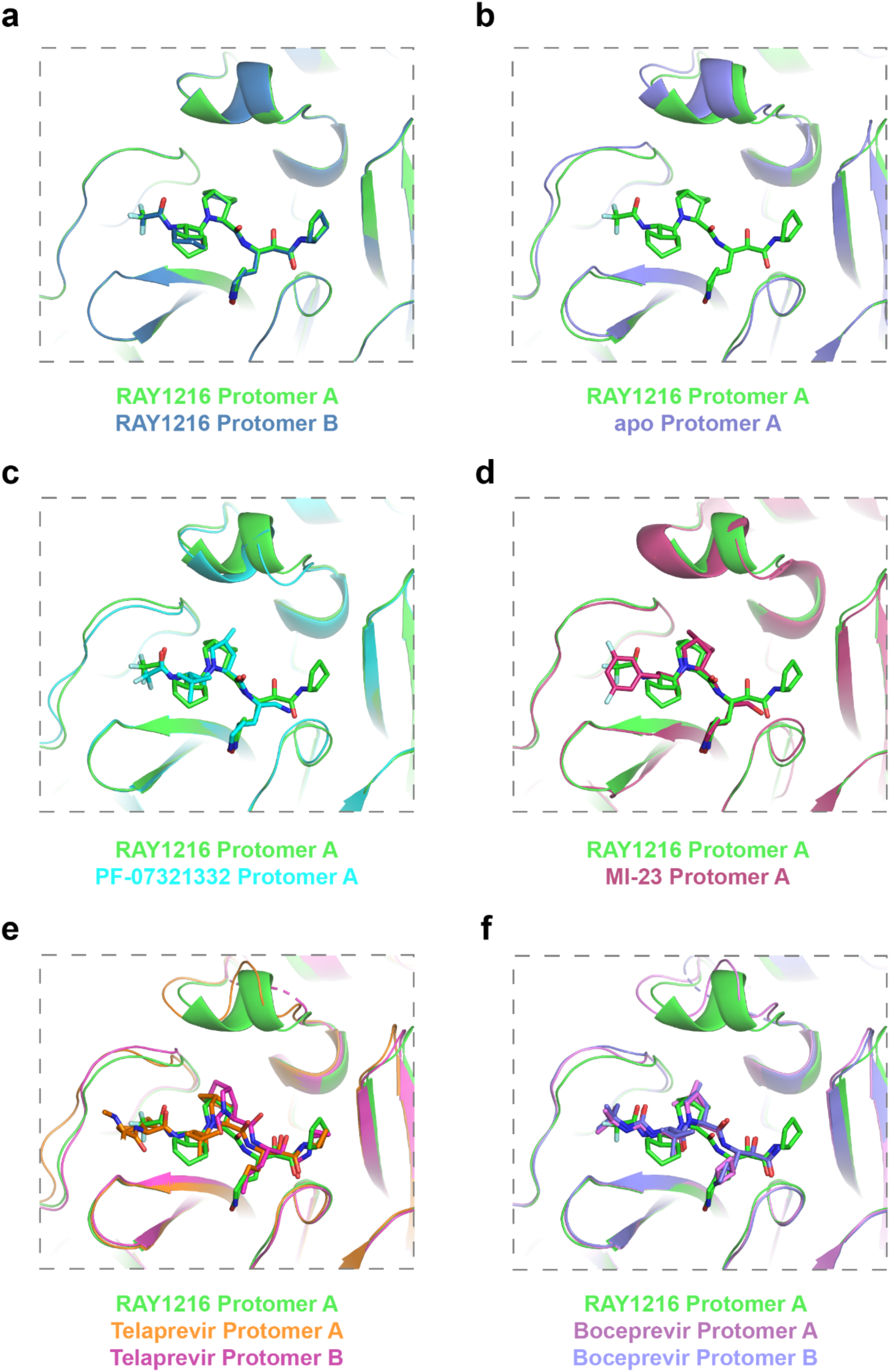
Comparison of active site structures of M^pro^ in different inhibitor complexes shows active site structural plasticity. Structures of boceprevir (PDB: 7com (Qiao *et al*., 2021)), MI-23 (PDB: 7d3i (Qiao *et al*., 2021)), PF-07321332 (PDB: 7rfw (Owen *et al*., 2021)), telaprevir (PDB: 7c7p (Qiao *et al*., 2021)) in complex with M^pro^ are compared with RAY1216:M^pro^ structures.

**Fig. S12.**
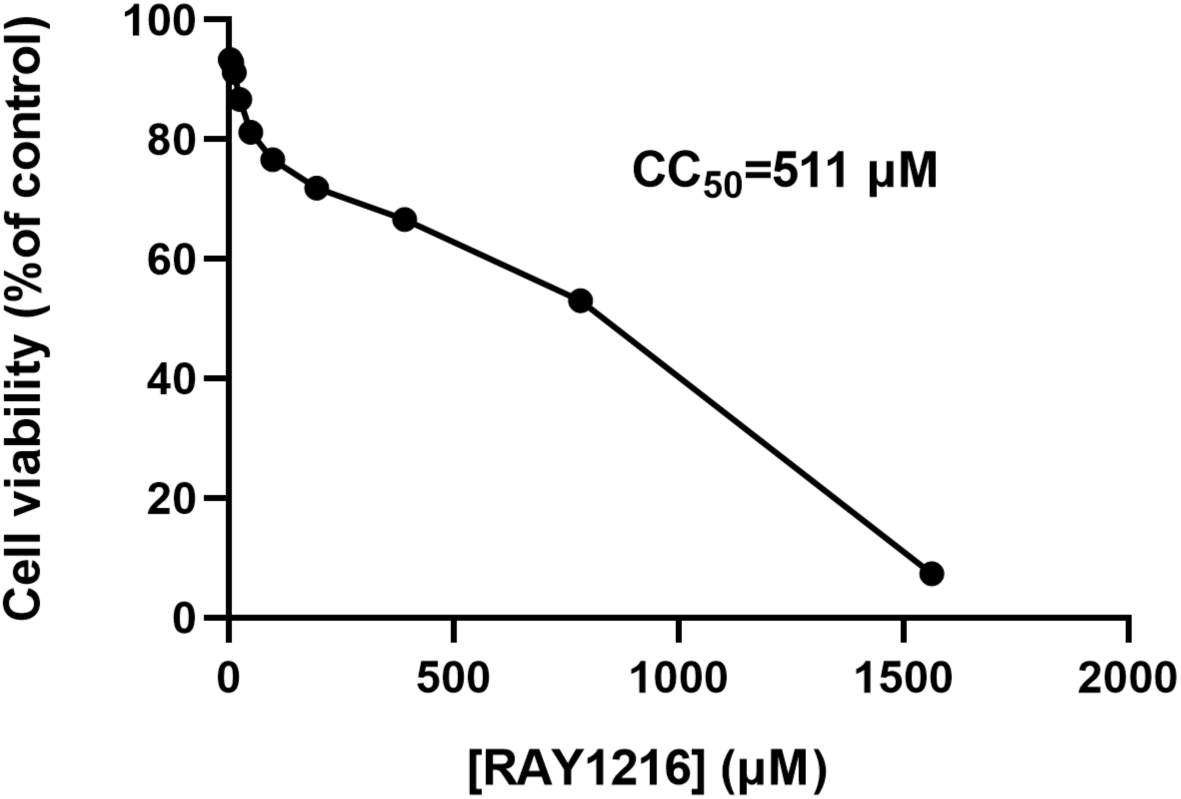
Cytotoxicity of RAY1216 on Vero E6 cells.

**Fig. S13.**
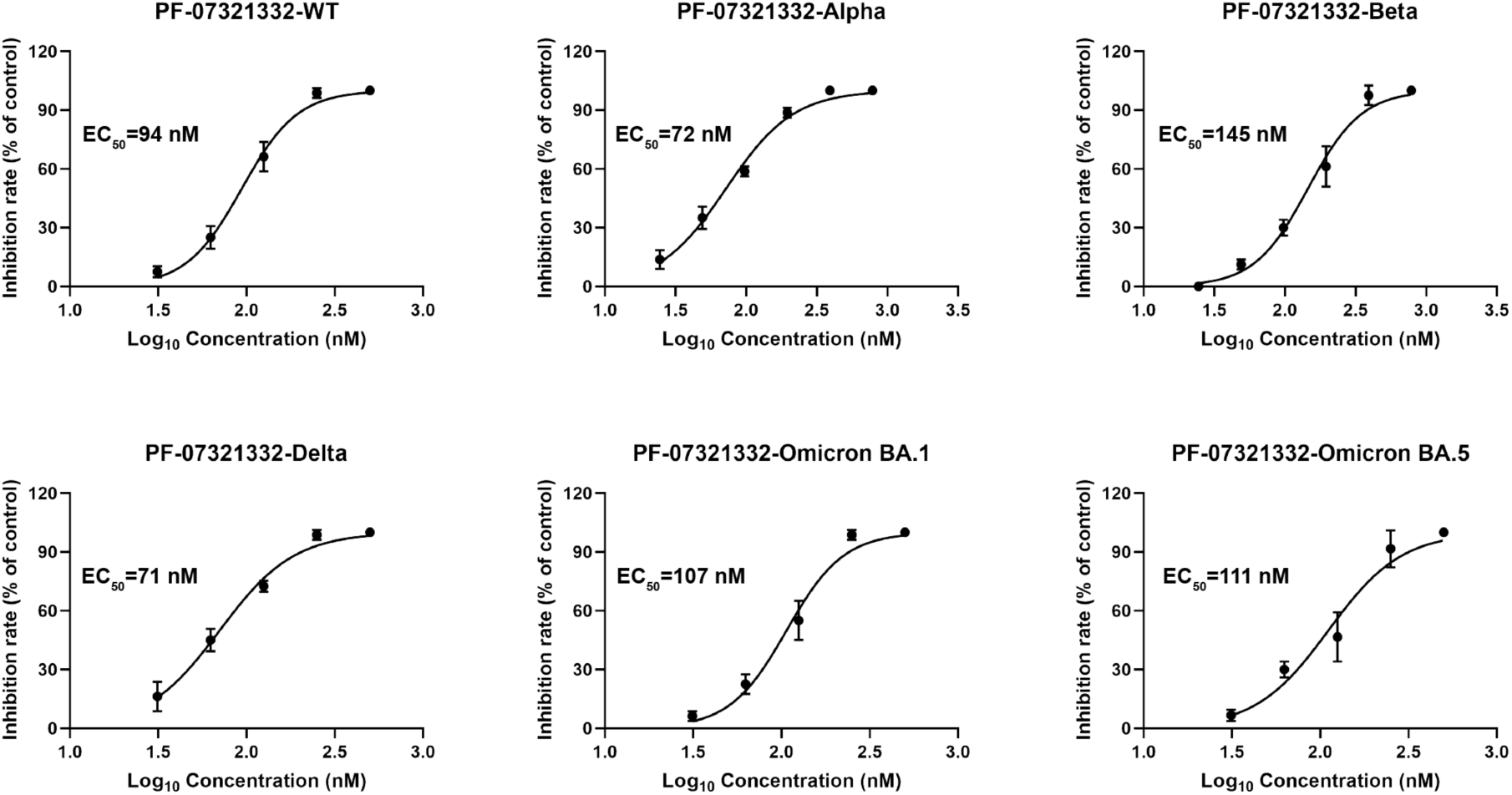
Dose–response curves (mean ± SD, *n* = 3) of wildtype (WT) and variant SARS-CoV-2 strains by PF-07321332 in Vero E6 cell using MTT cell viability assay.

**Table S6.** *In-vitro* plasma stability data.

| Animal species | Plasma stability (%2h) |  |
| --- | --- | --- |
|  | RAY1216 | PF-07321332 |
| mouse | 85 | 91.4 |
| rat | 85 | 84.3 |
| dog | 83 | 74.6 |
| cynomolgus macaque | 81 | 99.6 |
| human | 110 | 74.6 |

## Materials and Methods

### Synthesis of RAY1216

#### Step 1: Synthesis of the RAY1216-2 hydrochloride salt

To a solution of RAY1216-1 (500 mg, 1.75 mmol) in ethyl acetate (5 mL) at 20 °C HCl/EA (10 mL, 4 M) was added. The resulting mixture was stirred for 2 h at 20 °C. Solvent was removed to afford the residue as unpurified RAY1216-2 hydrochloride salt. ^1^H NMR (400 MHz, CD_3_OD) δ = 4.28 - 4.20 (m, 1H), 3.91 - 3.81 (m, 3H), 3.45 - 3.35 (m, 2H), 2.86 - 2.74 (m, 1H), 2.48 - 2.36 (m, 1H), 2.29 - 2.19 (m, 1H), 2.02 - 1.94 (m, 1H), 1.93 - 1.80 (m, 1H).

#### Step 2: Synthesis of RAY1216-4

To a solution of Boc-L-cyclohexylglycine (1 g, 3.89 mmol) in N,N-dimethylformamide (10 mL) 2-(7-azobenzotriazole)-N,N-tetramethylurea hexafluorophosphate (1.77 g, 4.66 mmol) was added. The resulting mixture was stirred for 0.5 h, to which diisopropylethylamine (1.26 g, 9.72 mmol) and RAY1216-3 hydrochloride salt (1.02 g, 4.66 mmol) were added. The resulting mixture was stirred for 16 h at 20 °C. The reaction mixture was added to methyl tert-butyl ether (50 mL), water (20 mL) and washed with 3% citric acid (20 mL×2) and brine (20 mL). The combined organic phase was dried over anhydrous sodium sulfate. Solvent was removed and the residue was purified by silica gel column chromatography (petroleum ether: ethyl acetate = 3:1) to afford RAY1216-4. ^1^H NMR (400MHz, CDCl_3_) δ = 5.22 - 5.11 (m, 1H), 4.36 (d, *J*=3.9 Hz, 1H), 4.27 (dd, *J*=6.9, 9.3 Hz, 1H), 4.21 - 4.12 (m, 2H), 3.83 (dd, *J*=7.8, 10.4 Hz, 1H), 3.70 (br dd, *J*=3.6, 10.4 Hz, 1H), 2.81 - 2.61 (m, 2H), 1.82 - 1.70 (m, 6H), 1.68 - 1.61 (m, 4H), 1.56 - 1.48 (m, 2H), 1.46 - 1.38 (m, 9H), 1.29 - 1.22 (m, 4H), 1.21 - 0.98 (m, 4H).

#### Step 3: Synthesis of RAY1216-5

To a THF (14 mL) solution of RAY1216-4 (1.41g, 3.34 mmol), LiOH•H_2_O (280.03 mg, 6.67 mmol) in water (5 mL) was added. The resulting mixture was stirred for 16 h at 20 °C. Citric acid was added to the reaction mixture to 3%. Solvent was removed and residue was extracted with ethyl acetate (50 mL), washed with brine (30 mL). The combined organic phase was dried over anhydrous sodium sulfate. Solvent was removed to afford the residue as unpurified RAY1216-5. ^1^H NMR (400MHz, DMSO-d_6_) δ = 12.58 - 12.23 (m, 1H), 6.92 - 6.82 (m, 1H), 4.11 - 3.94 (m, 2H), 3.82 - 3.76 (m, 1H), 3.72 - 3.62 (m, 1H), 2.73 - 2.64 (m, 1H), 2.62 - 2.55 (m, 1H), 1.92 - 1.42 (m, 12H), 1.40 - 1.32 (m, 9H), 1.18 - 1.06 (m, 3H), 1.00 - 0.81 (m, 2H).

#### Step 4: Synthesis of RAY1216-6

To a 2-butanone (7 mL) solution of RAY1216-5 (650 mg, 1.65 mmol), 1-hydroxybenzotriazole (222.63 mg, 1.65 mmol), 1-(3-dimethylaminopropyl)-3-ethylcarbodiimide hydrochloride (379.03 mg, 1.98 mmol), diisopropylethylamine (638.84 mg, 4.94 mmol) were added. The resulting mixture was stirred for 0.5 h at 20 °C, before RAY1216-2 hydrochloride salt (366.88 mg, 1.65 mmol) was added. The resulting mixture was stirred for 16 h at 20 °C. The reaction mixture was diluted with water (20 mL) and extracted with dichloromethane: methanol (30 mL×2, 10:1), the combined organic phase was washed with 3% citric acid (20 mL×2), brine (30 mL). The organic phase was dried over anhydrous sodium sulfate. Solvent was removed and the residue was purified by silica gel column chromatography (dichloromethane: methanol = 20: 1) to afford RAY1216-6. ^1^H NMR (400 MHz, CDCl_3_) δ = 7.49 - 7.42 (m, 1H), 6.23 - 6.05 (m, 1H), 5.28 - 5.17 (m, 1H), 4.64 - 4.51 (m, 1H), 4.43 - 4.24 (m, 2H), 3.92 - 3.81 (m, 1H), 3.78 - 3.70 (m, 3H), 3.39 - 3.27 (m, 2H), 2.94 - 2.75 (m, 2H), 2.57 - 2.36 (m, 2H), 2.24 - 2.07 (m, 1H), 1.94 - 1.50 (m, 14H), 1.49 - 1.41 (m, 9H), 1.27 - 0.95 (m, 6H).

#### Step 5: Synthesis of RAY1216-7

To a THF (31 mL) solution of RAY1216-6 (3.10 g, 5.51 mmol) at 0 °C, lithium borohydride (240.02 mg, 11.02 mmol) was added. Temperature of the mixture was allowed to warm to 20 °C slowly and stirred for 2 h at 20 °C. The mixture was diluted with water (10 mL) and ethyl acetate (20 mL) and stirred for 10 min, white solid was collected as crude target product RAY1216-7 by filtration. LCMS (m/z) 535.4 [M+1]^+^.

#### Step 6: Synthesis of RAY1216-8

To a DCM (10 mL) solution of compound RAY1216-7 (0.5 g, 935.13 μmol) Dess-Martin periodinane (594.94 mg, 1.40 mmol) was added. The resulting mixture was stirred for 16 h. The mixture was diluted with saturated sodium thiosulfate (15 mL) and saturated sodium bicarbonate solution (15 mL) and stirred for 10 min. The aqueous layer was extracted with DCM (50 mL×2). The combined organic phase was washed with brine (5 mL) and dried over anhydrous sodium sulfate. Solvent was removed and to afford the residue as unpurified RAY1216-8. LCMS (*m/z*) 533.4 [M+1]^+^.

#### Step 7: Synthesis of RAY1216-9

To a DCM (10 mL) solution of RAY1216-8 (436 mg, 818.52 μmol), acetic acid (58.98 mg, 982.22 mmol) and cyclopentyl isocyanide (94.44 mg, 982.22 μmol) were added. The resulting mixture was stirred for 2 h at 25 °C. After addition of saturated ammonium chloride solution (10 mL), the reaction mixture was stirred for 10 min and extracted with DCM (20 mL). The combined organic phase was washed with brine (5 mL). The organic phase was dried over anhydrous sodium sulfate, solvent was removed and the residue was purified by silica gel column chromatography (dichloromethane: methanol = 10:1) to afford RAY1216-9. LCMS (*m/z*) 688.4 [M+1]^+^.

#### Step 8: Synthesis of RAY1216-10

To a MeOH (3 mL) solution of RAY1216-9 (190 mg, 276.22 μmol), K_2_CO_3_ (95.44 mg, 690.54 μmol) in water (5 mL) was added. The resulting mixture was stirred for 16 h at 20 °C. The reaction mixture was diluted with 3% citric acid and extracted with DCM (40 mL× 3), the combined organic phase was washed with brine (30 mL) and dried over anhydrous sodium sulfate. Solvent was removed to afford the residue as unpurified crude RAY1216-10.

#### Step 9: Synthesis of RAY1216-11

To a DCM (10 mL) solution of RAY1216-10 (238.00 mg, 368.52 μmol), Dess-Martin periodinane (203.19 mg, 479.08 μmol) was added. The resulting mixture was stirred for 16 h at 20℃. The reaction mixture was diluted with saturated sodium thiosulfate (15 mL) and saturated sodium bicarbonate solution (15 mL), extracted with DCM (50 mL×2), and washed with brine (15 mL). The organic phase was dried over anhydrous sodium sulfate, solvent was removed, and the residue was purified by silica gel column chromatography (dichloromethane: methanol = 20:1) to afford RAY1216-11.

#### Step 10: Synthesis of RAY1216-12

To a THF (3 mL) solution of RAY1216-11 (125 mg, 194.16 μmol), HCl/EA (2.91 mL, 4 M) was added. The resulting mixture was stirred for 1 h at 25 °C. Solvent was removed to afford crude RAY1216-12.

#### Step 11: Synthesis of RAY1216

To a THF (3 mL) solution of RAY1216-12 (125 mg, 229.91 μmol) at 0 °C, TFAA (193.15 mg, 919.63 μmol) and pyridine (127.30 mg, 1.61 mmol) were added. The resulting mixture was stirred for 16 h at 20 °C. The reaction mixture was diluted with water (20 mL) and extracted with dichloromethane (30 mL× 2). The combined organic phase was washed with 3% citric acid (40 mL) and brine (40 m L×2). The organic phase was dried over anhydrous sodium sulfate. Solvent was removed and the residue was purified by HPLC to afford RAY1216. LC-MS (m/z) 640.0 [M+1]^+^.

^1^H NMR (400 MHz, DMSO-d6) δ ppm 9.75 d (J=7.5 Hz,1H) 8.64 d (J=7.5 Hz,1H), 8.50 d (J=8.3 Hz,1H), 7.65 s (1H), 5.14 ddd (J=11.5, 8.3, 2.9 Hz,1H), 4.28 t (J=8.6 Hz,1H), 4.20 d (J=4.2 Hz,1H), 4.04 m (1H), 3.76 AABB-d (J=10.2, 7.6, 3.0 Hz,2H), 3.20 - 3.11 m (2H), 2.69 m (1H), 2.55 m (1H), 2.50 m (1H), 2.20 - 1.69 m (2H), 1.88 - 1.60 m (2H), 1.83 m (1H), 1.80 - 1.40 m (6H), 1.80 - 1.50 m (4H), 1.80-1.40 m (4H), 1.79-0.95 m (4H), 1.65 - 1.13 m (6H). ^13^C NMR (400 MHz, DMSO-d6) δ ppm 115.9, 156.5, 56.1, 38.7, 28.2, 28.6, 25.4, 25.6, 25.8, 168.6, 53.5, 42.7, 24.6, 31.2, 31.66, 47.5, 65.5, 171.9, 51.8, 31.8, 37.5, 178.4, 39.5, 27.5, 197.2, 161.0, 50.4, 23.6, 31.64, 31.68.

### Crystallization of RAY1216 and X-ray diffraction

RAY1216 powder (approximately 30 mg) was dissolved in isopropyl acetate (approximately 600 μl) with gentle stirring gently until the mixture was saturated. The solution was transferred into a transparent 2ml MS sample vial after being filtered by syringe filter. The sample vial was sealed with a parafilm and 5-10 holes on the parafilm were pierced by a syringe needle. The MS vial was placed in a closed brown bottle which contained 0.1 - 0.2 cm level of n-hexane. Solutions were let stand at 20 - 30 °C for 48 hours. After granular white crystals were observed in the MS vial, the isopropyl acetate was removed, and a single crystal of RAY1216 was sealed with silicone grease and subjected to X-ray diffraction on a Rigaku Oxford Diffraction XtaLAB Synergy-S four-circle diffractometer equipped with a CuKα source (λ=1.54184 Å) and a HyPix-6000HE area detector.

### Data collection and structure determination

40929 diffraction spots were collected by X-ray diffraction experiment, and 5638 independent diffraction dots were indexed and integrated (R_int_=0.0391). Diffraction collection range 2θ = 5.752° to 133.106°, diffraction index range −12≤h≤12, −11≤k≤11, −18≤l≤18. Structure was determined by SHELXT (Sheldrick, 2015b) and refined by SHELXL (against F^2^) (Sheldrick, 2015a). 406 parameters are participating in structural refinement. The final result has a goodness-of-fit (s) = 1.041, R_1_ = 0.0351, wR_2_ = 0.0914. The residual electron density values are 0.39 and −0.31 e Å^−3^. The data collection and structure refinement statistics are summarized in **Table S1.**

### Recombinant protein production

Based on a previous study (Zhang *et al*., 2020), a construct encoding SARS-CoV-2 Mpro (ORF1ab 3264-3569, GenBank code:MN908947.3) was subcloned into the pGEX-6p-1 vector between the BamHI and XhoI restriction sites with extra C-terminal extension GPHHHHHHHHHH. The resulting construct was verified by DNA sequencing. The construct plasmid was transformed into BL21 (DE3) *E. coli* cells (Vazyme, #C504-02/03) and scale-up expression (∼ 6 L) was started from a single colony in LB medium supplemented with 100 μg/ml of ampicillin at 37 °C. The cells were induced with 0.5 mM IPTG when the OD600 reached 0.8. Cells were allowed to grow post induction for 20 h at 16 °C.

The cells were harvested by centrifugation, and cell pellet was lysed in the lysis buffer (20mM Tris, pH 7.8, 150mM NaCl, 10mM imidazole) by sonication on ice. Cell lysate was cleared by high-speed centrifugation (20,000 × g at 4°C for 1h). The supernatant was mixed with Ni-NTA resin for ∼ 2 h at 4 °C on a shaker. The Ni-NTA resin was washed with two buffers of different imidazole concentrations (20 mM Tris, pH 7.8, 150 mM NaCl, 20 mM/50 mM imidazole) each for over 30 resin volumes to remove contaminants. The target protein was eluted by the elution buffer (20 mM Tris, pH 7.8, 150 mM NaCl, 500 mM imidazole). 400U Human rhinovirus (HRV) 3C enzyme was added into the eluted protein to remove the C-terminal histidine tag and the mixture was dialysed at 4 °C overnight in dialysis buffer (20 mM Tris, pH 7.8, 150 mM NaCl, 1m M DTT) using a dialysis bag with MWCO (Molecular Weight Cut Off) of 10 kDa. The dialysed mixture was reloaded onto the Ni-NTA resin and His-tag-free target protein was collected from the flow-through.

Since the expressed M^pro^ contains the native M^pro^ cleavage sequence “SAVLQ/SGFRK” found between Nsp4 and Nsp5 (M^pro^) in the SARS-CoV-2 Nsp polyprotein (slash indicates the M^pro^ cleavage site) near the N-terminus, M^pro^ auto-cleaving activity generates an authentic N-terminus during protein expression. The HRV 3C enzyme recognition site has the sequence (LEVLFQ/GP, slash indicates the HRV 3C enzyme cleavage site) after HRV 3C cleavage, it will generate an authentic M^pro^ C-terminus (LEVLFQ/SAV, native M^pro^ C-terminal recognition sequence, slash indicates the cleavage site). Purified M^pro^ with authentic N- and C-termini was concentrated by a 10 kDa MWCO Amicon Ultra 50 centrifugal filters (Merck Millipore) at 4 °C to ∼ 10 mg/ml. Concentrated protein was either used for crystallization without freezing or flash frozen in liquid nitrogen and stored under −80 °C.

### Enzyme activity assay

The enzyme assays were performed in enzyme kinetics assay buffer (20 mM Tris pH 7.8, 150 mM NaCl, 1 mM DTT and 100 μg/ml bovine serum albumin) using Dabcyl-KTSAVLQ/SGFRKME-Edans (Beyotime, #P9733, ‘/’ indicates the M^pro^ cleavage site) as the substrate. Florescent signal by enzyme cleavage of the substrate was monitored on a Molecular Devices FlexStation 3 reader with filters for excitation at 340 nm and emission at 490 nm at 20 °C.

### Molar response coefficient of fluorescent product

M^pro^ at a relatively high concentration (1.0 μM) was assayed with 1.25, 2.5, 5, 10, 20, 40, and 80 μM substrate at 37 °C. Under these experimental conditions the substrate was completely converted into the fluorogenic product over a course of 20 minutes. The total observed change in fluorescence intensity (Δ*F*), when plotted against the starting concentration of substrate ([S]), displayed a significant involvement of the frequently seen Dabcyl-Edans inner-filter effect (Liu et al., 1999). The dependence of Δ*F* on [S] follows a non-linear quadratic function as illustrated in **Fig. S6a**. However, within the restricted range of substrate concentrations (1.25 - 20 μM, **Fig. S6b**) the dependence of Δ*F* on [S] is approximately linear, with the slope equal to the differential molar response coefficient of the product *r*_P_ = (88.1 ± 3.5) rfu/μM. Thus, in all the following enzyme kinetic analyses we constrained the substrate concentration accordingly ([S] ≤ 20 μM) and treated the molar response coefficient as a fixed parameter, set to the best-fit value of *r*_P_ = 88 rfu/μM.

### Determination of the Michaelis constant, *K*_M_

Preliminary attempts to determine the Michaelis constant *K*_M_ by fitting initial rates vs [S] to the Michaelis-Menten equation were found to be inaccurate due to the Dabcyl-Edans inner-filter effect (Liu *et al*., 1999). Thus, substrate kinetic parameters were determined by the global fit of reaction progress curves recorded at 1.25, 2.5, 5, 10, and 20 μM to the first-order ordinary differential-equation (ODE) model corresponding to the reaction mechanism shown in **Scheme 1**, using the software package DynaFit (Kuzmic, 1996; 2009). The DynaFit script file is provided in the Supplementary Dataset.

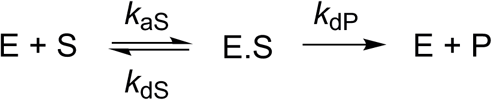

#### Scheme 1. Assumed Michaelis-Menten reaction mechanism of M^pro^ substrate hydrolysis

The mathematical model for the reaction progress curves is auto-generated by DynaFit according to Scheme 1 and is shown as **Eq. 1**, where *F* is the fluorescence intensity at the arbitrary reaction time *t*; *F*_0_ is the baseline offset or fluorescence intensity observed at time *t* = 0; *r*_P_ = 88 rfu/μM is the molar response of the reaction product; and [P] is the concentration of the reaction product at time *t*. In its turn, the product concentration is computed by numerically solving the ODE system represented by **Eq.1–Eq.5**.

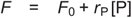

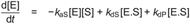

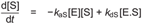

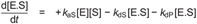

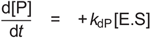

The enzyme–substrate association rate constant *k*_a_S was fixed at the diffusion limited constant value of 1.0 μM^−1^s^−1^ (or 1 × 10^6^ M^−1^s^−1^) (Fersht, 1999), whereas the dissociation rate constants *k*_d_S and *k*_d_P were treated as globally adjustable model parameters. Each individual progress curve was associated with a locally optimized offset parameter *F*_0_. The experimental data files (Km-R1-f.csv and Km-R2-f.csv) are provided as **Supplementary Datasets**. The results of fit for both replicates are illustrated graphically in **Fig. S7**.

The best-fit values of adjustable model parameters are listed in **Table S2**. The average and standard deviation from replicates (n = 2) of the dissociation rate constant is *k*_dS_ = (31.0 ± 0.2) s^−1^. This particular value of *k*_dS_ is utilized in subsequent kinetic analyses of M^pro^ inhibition as a fixed model parameter (see below). The corresponding average and standard deviation of the Michaelis constant is *K*_M_ ≡ (*k*_dS_ + *k*_dP_)/*k*_aS_ = (31.0 ± 0.2) μM. This value is identical, within the specified experimental error, to a previously published value of *K*_M_ = (28.2 ± 3.4) μM (Ma *et al*., 2020).

### Inhibition kinetics of PF-07321332 and active-site titration

The microscopic rates constants for association and dissociation of PF-07321332, as well as, the concentration of M^pro^ active sites hence the enzyme’s turnover number *k*_cat_, were determined as follows. The enzyme (nominal concentration 80 nM) was assayed at varied concentrations of the inhibitor (maximum 444 nM, 2/3 dilution series down to 17 nM, nine concentrations plus control [I] = 0) in triplicate. The ten reaction progress curves from each replicate were combined into a global dataset and fit to a differential-equation model corresponding to the reaction mechanism shown in **Scheme 2** using the software package DynaFit (Kuzmic, 1996; 2009). The requisite DynaFit script file and PF-07321332 inhibition progress curve data are provided as **Supplementary Datasets**.

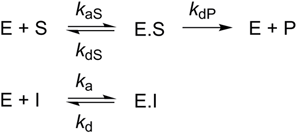

#### Scheme 2. Proposed reaction and inhibition mechanisms of M^pro^

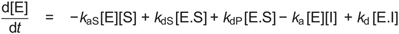

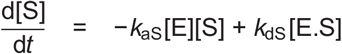

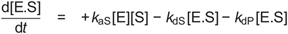

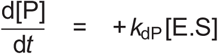

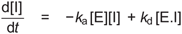

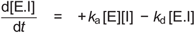

To obtain kinetic parameters of PF-07321332 inhibition, the microscopic rate constants *k*_aS_ and *k*_dS_ are fixed, whereas the turnover number rate constant *k*_cat_ ≡ *k*_dP_ and the active enzyme site concentration are treated as adjustable model parameters (**Table S3**) (marked by the notation “??” in DynaFit script files). The combined progress curves were fit globally to **Eq.1**, where the product concentration [P] in this case is computed by numerically solving the ODE system represented by **Eq.6–Eq.11**.

Thus, the complete list of globally optimized parameters (shared by all progress curves) consists of *k*_dP_, *k*_a_, *k*_d_, and [E]; the ten baseline offset values, *F*_0_, each specific to a particular progress curve, were treated as locally adjustable model parameters. The overlay of experimental data on the best-fit model curves and the corresponding plots of instantaneous reaction rates are shown in **Fig. S8**. Numerical results are summarized in **Table S3**.

The results listed in **Table S3** show that all adjustable model parameters were obtained with better than 11% reproducibility. The enzyme was approximately 61% active. The turnover number *k*_cat_ = 0.12 s^−1^ compares well with *k*_cat_ = 0.16 s^−1^ previously reported (Ma *et al*., 2020). Similarly, the inhibition constant *K*_i_ ≡ *k*_d_/*k*_a_ = (3.8 ± 0.2) nM determined here compares well with the previously reported value of *K*_i_ = 3.1 (1.5 − 6.8) nM (Owen *et al*., 2021). The PF-07321332 inhibitor associates relatively rapidly with the enzyme, with bimolecular association rate constant equal to *k*_a_ = 4.9 × 10^5^ M^−1^s^−1^. The drug-target residence time *t*_res_ ≡ 1/*k*_d_ is approximately 9 minutes.

### Inhibition kinetics of RAY1216

Preliminary analysis of RAY1216 inhibition data revealed that it is possible to reliably determine either the turnover number *k*_cat_ ≡ *k*_dP_, or the active enzyme concentration [E], but not both. However, the precise values of these model parameters and in particular the active enzyme concentration strongly influence the best-fit values of the inhibition rate constants *k*_a_ and *k*_d_. For this reason we have analysed the RAY1216 datasets while treating the microscopic rate constant *k*_dP_ as a fixed parameter determined in the fit of the PF-07321332 data above (The RAY1216 and PF-07321332 experiments were carried out in parallel on the same day using the same M^pro^ prep). (The “??” notation is deleted on the “`kdP`” line in the provided DynaFit script to fix “*k*_dP_” for fitting the RAY1216 datasets.)

Thus, the complete list of globally optimized parameters (shared by all progress curves) consists of *k*_a_, *k*_d_, and [E]; the ten baseline offset values, *F*_0_, each specific to a particular progress curve, were again treated as locally adjustable model parameters. The overlay of experimental data on the best-fit model curves and the corresponding plots of instantaneous reaction rates are shown in **Fig. S9**. Note that, in comparison with **Fig. S8**, the instantaneous rate plots show significantly longer time that is required for near-equilibrium to be established, even at the highest inhibitor concentrations. This graphically illustrates that RAY1216 is a “slow-on, slow-off” inhibitor of M^pro^, while PF-07321332 is a “fast-on, fast-off” inhibitor. Numerical results of fit are summarized in **Table S4**.

The results listed in **Table S4** show that all adjustable model parameters were obtained with better than 5% reproducibility. The enzyme was apparently 66% active. This value differs slightly from the 61% enzyme activity determined in the analysis of PF-07321332 data (see **Table S3**). We observed that, throughout the working day, the enzymatic activity of M^pro^ decreased slightly but systematically over time. RAY1216 associates relatively slowly with the target enzyme, with bimolecular association rate constant equal to *k*_a_ = 1.9 × 10^4^ M^−1^s^−1^ The inhibition constant of RAY1216, *K*_i_ = 8.4 nM, is 2.2 times higher than the *K*_i_ for PF-07321332 determined here (**Table S3**). In contrast, the drug-target residence time is 104/9 = 11.5 times longer.

### Crystallization and crystal soaking with RAY1216 inhibitor

Apo M^pro^ crystals were crystallized by mixing 1 μl of freshly purified M^pro^ (without freezing) at 10 mg/ml with 1 μl crystallization solution (0.1 M MES monohydrate pH 6.5, 12% w/v Polyethylene glycol 20,000) using hanging drop vapor diffusion method at 16 °C.

Crystals normally grew overnight. The apo crystals were flash frozen in cryoprotection solution (0.1 M MES monohydrate pH 6.5, 12% w/v polyethylene glycol 20,000, 40% glycerol) using liquid nitrogen. To obtain RAY1216 soaked crystals, apo crystals were transferred into the crystallization solution supplemented with 6.6 mM RAY1216 and 3% DMSO (from the RAY1216 solution). The crystals were soaked for ∼10 min at 16 °C. Finally, crystals were briefly soaked in cryoprotection solution (0.1 M MES monohydrate pH 6.5, 12% w/v polyethylene glycol 20,000, 40% glycerol) supplemented with 6.6 mM RAY1216 before being frozen in liquid nitrogen.

### Data collection and structure determination

Single crystal X-ray diffraction data were collected on beamline BL19U1 at Shanghai Synchrotron Radiation Facility (SSRF) at 100 K using an Eiger X 16M hybrid-photon-counting (HPC) detector. Data integration and scaling were performed using the XDS software (BUILT 20220220) (Kabsch, 2010). Structures were determined by molecular replacement (MR) using the Phaser MR 2.8.3 (McCoy et al., 2007) program in CCP4 7.1.018 (Winn et al., 2011), with a SARS-CoV-2 M^pro^ structure (Zhao *et al*., 2022) (PDB code: 7VH8) as the search model. Iterative manual model building was carried out in Coot 0.9.6 (Emsley and Cowtan, 2004), Final structures were refined with Refmac 5.8.0267 (Murshudov et al., 1997). The data collection and structure refinement statistics are summarized in **Table S5**.

### Cell lines and virus strains

African green monkey kidney epithelial (Vero E6) cells were purchased from the American Type Culture Collection (ATCC), and cultured in Dulbecco’s modified Eagle’s medium (DMEM, Gibco, USA) supplemented with 10% fetal bovine serum (FBS, Gibco, USA), 100 μg/mL streptomycin (Gibco, USA), and 100 U/mL penicillin (Gibco, USA). SARS-CoV-2 and its variants, namely Alpha (B.1.1.7), Beta (B.1.1.529), Delta (B.1.617.2), Omicron BA.1 (B.1.1.529) and Omicron BA.5 (BA.5.2) were isolated from clinical samples and were deposited at the First Affiliated Hospital of Guangzhou Medical University. Viruses were propagated as previously described (Zhu et al., 2020a), the viruses were aliquoted and stored at −80 °C and the titres of cultured viruses were estimated as 50% tissue culture infective doses (TCID_50_) using the Reed–Muench method.

### Cytotoxicity and cytopathic effect (CPE) inhibition assays-SARS-CoV-2

The 50% toxicity concentration (CC_50_) for RAY1216 in Vero E6 cells was determined using the MTT (3-(4,5-dimethylthazolk-2-yl)-2,5-diphenyl tetrazolium bromide) assay (Park et al., 2011). Different dilutions of RAY1216 and PF-07321332 were incubated with Vero E6 (5 × 10^4^ cells/well) cells in 96-well plates for the cytotoxicity assay, and the concentration of RAY1216 and PF-07321332 causing 50% cell death were determined as the CC_50_ value. The 50% inhibition concentration (EC_50_) of virus-induced cytopathic effect (CPE) was used to investigate the efficacy of RAY1216 and PF07321332 against SARS-CoV-2. A monolayer of Vero E6 cells was inoculated with 100 TCID_50_ of SARS-CoV-2 wildtype or variant strains at 37 °C for 2 h. The cells were incubated with different concentrations of RAY1216 and PF-07321332 after removal of the inoculum. Infected cells were observed under a microscope after 72 h of incubation to assess CPE. Dose response curves were plotted as CPE vs Log inhibitor concentrations. The selectivity indices (SI) were determined using the ratio of CC_50_ to EC_50_.

### Antiviral and anti-inflammatory activity of RAY1216 in mouse model

Antiviral studies using animals were approved by the Guangzhou Medical University Ethics Committee of Animal Experiments (IACUC certificate No.: GZL0008). All antiviral experiments using animals passed the ethical review and were performed in strict accordance with the National Research Council Criteria and the Chinese Animal Protection Act. Five-to six-week-old female ACE2 transgenic C57BL/6 mice (Bao *et al*., 2020; Ma et al., 2022) weighing 18–22 g were acquired from GemPharmatech Co., Ltd. (Jiangsu, China) and housed under specific pathogen-free (SPF) conditions at Guangzhou Customs District Technology Center Biosafety Level 3 (BSL-3) Laboratory. Mice were randomly divided into six groups (*n* = 7): the control group; SARS-CoV-2 virus (Delta variant (B.1.617.2)) infected group; treatment groups of three different RAY1216 concentrations (600 mg/kg/day, 300 mg/kg/day, 150 mg/kg/day); and a PF-07321332 treatment group (600 mg/kg/day). Mice were anesthetized by inhalation of 5% isoflurane and each mouse was inoculated with 50 μl PBS containing a lethal dose of 10^5^ PFU SARS-CoV-2 (Delta variant) for the infected groups. For the control group, 50 µl PBS was administered intranasally. 2 hr after infection, the infected mice were intragastrically administered with RAY1216 (600 mg/kg/day, 300 mg/kg/day, 150 mg/kg/day), PF-07321332 (600 mg/kg/day) or PBS daily for 5 days. Weight change and mortality of mice in each group was recorded daily for 5 days. To measure lung virus titres and to examine lung pathology, a separate set of experiments was performed under the same grouping and conditions except that each mouse was inoculated with a non-lethal dose of 10^3.5^ PFU SARS-CoV-2 (Delta variant) for the infected groups. At 3 and 5 days post infection, mice were sacrificed, and lung tissues were collected to measure virus titres and to examine lung pathology.

### *In-vitro* plasma stability analysis

To assess *in-vitro* plasma stabilities of RAY1216 and PF-07321332, 2 µM of the tested compounds were incubated in plasma from different CD-1 mouse, SD rat, beagle dog, cynomolgus monkey and human at 37 °C for 2 hr. 40 μL of samples were added into 160 μL of internal standard working solution (200 ng/mL tolbutamide in MeOH). The mixed solutions were vortex and centrifuged at 16000 g for 10 min at 2-8 °C. The supernatant was analysed by LC-MS/MS.

### Pharmacokinetic studies

The pharmacokinetic (PK) studies using animals were approved by the WuXi AppTec Ethics Committee of Animal Experiments (IACUC certificate No.: NJ-20220531). All animals used in this study were male and chosen randomly. Pharmacokinetic properties of RAY1216 and PF-07321332 following a single intravenous injection (IV) or gavage (PO) administration were examined. Briefly, two groups were assigned for each animal species, male CD-1 mouse (2 mice/group), male SD rat (3 rats/group) and male cynomolgus macaque (2 cynomolgus macaques /group), respectively. The test compound was administered to each group at the indicated dose, orally or intravenously. The specific doses administered are shown in **Table 3**. Plasma samples of animals in IV groups were collected before administration (0), 0.083, 0.25, 0.5, 1, 2, 4, 8 and 24 hours after administration; samples of animals in PO groups were collected before administration (0), 0.25, 0.5, 1, 2, 4, 6, 8, and 24 hours after administration.

### LC-MS/MS analysis of plasma samples

Plasma samples were obtained by centrifugation and concentrations of the compounds in the serum were assessed by LC-MS/MS. On an ACQUITY UPLC System, ACQUITY UPLC Protein BEH C4 column (300Å, 1.7 μm, 2.1 × 50 mm), ACQUITY UPLC HSS T3 column (1.8 μm, 2.1 × 50 mm) and Phenomenex Kinetex C18 LC column (2.6 μm, 100 Å, 2.1 × 50 mm) were used for analyses of samples from mouse, rat and cynomolgus macaque, respectively. The pharmacokinetic parameters of RAY1216 and PF-07321332 in plasma were calculated using non-compartmental analysis as implemented in Phoenix WinNonlin software (version 8.3.4).

